# HLA micropolymorphisms confine neoantigen conformational adaptability and guide T cell receptor selectivity

**DOI:** 10.64898/2025.12.22.696040

**Authors:** Jiaqi Ma, Cory M. Ayres, Chad A. Brambley, Bassant Eldaly, W. W. J. Gihan Perera, James A. Lazar, Evgenii L. Kovrigin, Smita S. Chandran, Christopher A. Klebanoff, Brian M. Baker

**Affiliations:** Harper Cancer Research Institute and the Department of Chemistry and Biochemistry, University of Notre Dame, Notre Dame, IN USA; Human Oncology and Pathogenesis Program and the Parker Institute for Cancer Immunotherapy, Memorial Sloan Kettering Cancer Center (MSKCC), New York, NY, USA; Weill Cornell Medical College, Cornell University, New York, NY, USA

## Abstract

T cell receptor (TCR) restriction by highly polymorphic major histocompatibility complex (MHC) proteins is a foundation of cellular immunity. Although the effects of MHC polymorphisms on peptide binding and selection are well established, how micropolymorphisms within MHC supertypes impact immune recognition is poorly understood. Here, we identified a novel mechanism through which the micropolymorphisms in two closely related HLA-A3 superfamily members determine TCR specificity. We previously showed that TCRs specific for a public neoantigen from the *PIK3CA* oncogene presented by HLA-A*03:01 were unable to recognize the same epitope in the context of HLA-A*03:02. We found here that the micropolymorphisms distinguishing A*03:02 from A*03:01 exert their effect not by altering peptide binding or static structures, but by changing the conformational ensemble of the neoantigen in the groove, preventing it from adopting a conformation compatible with TCR binding. The effect is rooted in how the two polymorphic sites interact with other co-varying, evolutionarily coupled polymorphisms, reflecting a cross-groove network of interactions that controls the conformational adaptability of the peptide/HLA complex. We suggest polymorphism-dependent conformational adaptability reflects an evolved feature of class I MHC proteins that amplifies the impact of peptides in the groove, further diversifying epitopes and contributing to how TCRs and other immunoreceptors differentiate between antigens. Beyond this mechanistic insight, our findings emphasize the need for high-resolution HLA typing in efforts across immunology, including antigen-specific immunotherapy.

## Introduction

T cell receptor (TCR) recognition of antigenic peptides underlies cellular immunity, directing immune responses to pathogens and cancer as well as driving pathologies such as autoimmunity, transplant rejection, and graft vs. host disease. TCR recognition of peptide targets forms the basis of multiple efforts in immunotherapy, including the design and engineering of peptide-based vaccines, TCR-modified T cells, and soluble biologics ^1^. Central to both natural and engineered immunity is the phenomenon of MHC restriction, whereby TCRs only recognize peptides bound and presented by major histocompatibility complex proteins (HLA proteins in humans) ^2^. The genes encoding the classical MHC proteins are the most polymorphic in the human genome, with thousands of variants distributed throughout populations ^3^. Human HLA variants are grouped into 9-12 supertypes, defined primarily by their peptide binding preferences and distinguished by up to dozens of amino acid substitutions ^4^. Within each supertype are numerous variants that can differ by as few as 1-2 amino acids, commonly referred to as micropolymorphisms.

How class I MHC supertypes differentially bind peptides and the role this plays in T cell specificity and the overall diversification of the immune response is generally well understood. Less understood are the impacts of micropolymorphisms between closely related alleles within supertypes. While often assumed to play less significant roles, there are well known instances where micropolymorphisms have dramatic immune consequences. In autoimmunity, for example, variants of HLA-B*27, HLA-B*39, HLA-B*51 are differentially associated with ankylosing spondylitis, type 1 diabetes, or Beçhet’s syndrome ^5–7^. In the immune response to HIV, variants of HLA-B*57 are differentially associated with elite control of infection ^8,9^. Drug hypersensitivity has in some cases been shown to correlate with select variants ^10^, and outcomes in transplantation can be strongly impacted by subtle class I MHC mismatches ^11,12^. The molecular bases of these findings have been examined in many studies. In some cases, micropolymorphisms that distinguish closely related alleles can alter peptide binding and lead to structurally divergent peptide conformations ^13–19^, although this can vary with the specific peptides or alleles involved ^15,20–23^. Micropolymorphisms can also influence the molecular motions of peptides within the peptide binding groove ^22,24–27^, as well as the motions and structural stability of the groove itself ^28–32^.

We recently characterized T cell recognition of a public cancer neoantigen (neoAg; sequence ALHGGWTTK) derived from phosphoinositide 3-kinase α (PI3Kα), one of the most frequently mutated proteins in human cancer ^33^. We generated a panel of TCRs restricted to this epitope presented by HLA-A*03:01 (referred to as A*03:01 throughout). These TCRs, however, failed to recognize the neoAg in the context of HLA-A*03:02 (referred to as A*03:02), despite the peptide being presented equally well by both alleles. We initially hypothesized that the specificity for A*03:01 over A*03:02 could arise from conformational changes caused by the two amino acid differences that distinguish the alleles, found in positions 152 and 156 near the center of the α2 helix (Glu152 and Leu156 in A*03:01; Val152 and Gln156 in A*03:02).

Subsequently though, we showed that TCR recognition of the PI3Kα neoAg presented by A*03:01 requires a dramatic structural rearrangement in the peptide, among the largest seen in TCR binding of peptides presented by class I MHC proteins ^34^. The rearrangement, consisting of a flip in the center of the peptide, is impacted by the identity of the peptide position 2 anchor: a change in the anchor alters molecular motions in the base of the binding groove, influencing the ability of the peptide to move into a conformation compatible with TCR binding. This allosteric effect occurs without changes to the static structure of the unbound (apo) neoAg/A*03:01 complex, but was evident via spectroscopic and computational experiments that revealed anchor-dependent shifts in the dynamic ensemble that surrounds the average, crystallographic structure ^34^.

With this insight, we hypothesized that TCR discrimination between the PI3Kα neoAg presented by A*03:01 and A*03:02 could emerge from an allele-dependent impact on peptide and protein dynamics and thus conformational adaptability. Investigating this hypothesis, we found that the micropolymorphisms distinguishing the two alleles indeed alter the conformational ensemble of the peptide and, in A*03:02, prevent it from adopting the conformation required for specific TCR binding. Notably though, as with peptide binding affinity, the static structures of the neoAg/A*03:01 and neoAg/A*03:02 complexes are indistinguishable. Further, either of the two polymorphisms in A*03:01 alone, which individually define the rare alleles HLA-A*03:07 and HLA-A*03:65 (referred to as A*03:07 and A*03:65), were sufficient to ablate TCR recognition. These outcomes arise from how the polymorphisms alter electrostatic interactions with other co-varying, evolutionarily coupled polymorphic positions that span the peptide binding groove. In addition to the neoAg/A*03:01 complex, we identified this cross-groove connection in structures of other peptides bound to A*03:01, but as we found with the neoAg bound to A*03:02, A*03:07, and A*03:65, it is altered or non-existent with other members of the HLA-A3 superfamily.

Overall, our results reveal the evolution of a sophisticated mechanism for diversifying how peptides are perceived within the context of class I MHC proteins. They intersect with recent work showing how conformational flexibility within peptide/MHC binding grooves can dramatically influence immunological outcomes, even in the absence of static structural differences ^35–39^. This last point is particularly relevant given the growing interest in peptide/MHC structural modeling and prediction, emphasizing the need to consider not just static structures but conformational ensembles and how they can be allosterically tuned in efforts to predict immunogenicity from structural features ^40–43^. Lastly, our findings expand the range of functional consequences attributable to micropolymorphisms and reinforce the need to consider high resolution HLA typing in efforts ranging from immunotherapy to transplantation, as the same epitopes presented by closely related molecules can clearly be perceived as different antigens.

## Results

### PI3Kα neoAg/HLA-A*03-specific TCRs distinguish between A*03:01 and A*03:02

Our previous work showed that T cells expressing TCRs specific for the PI3Kα neoAg presented by A*03:01 failed to recognize the same peptide when presented by A*03:02, even though the two complexes were of identical stability ^34^. A*03:01 differs from A*03:02 by two micropolymorphisms: E152 and L156 in A*03:01, and V152 and Q156 in A*03:02. Both positions are in the α2 helix, after the linker that connects the short and long arms of the helix and adjacent to the C-terminal half of the peptide (**Fig. 1A**). The two micropolymorphisms also exist individually in human populations, albeit at very low frequencies: E152V alone is A*03:07, with an overall frequency of 0.000074, and L156Q alone is A*03:65, with a frequency of 0.000002 (for comparison, A*03:01 is present at a frequency of 0.151, and A*03:02 is present at a frequency of 0.0086) ^44^.

**Figure 1.**
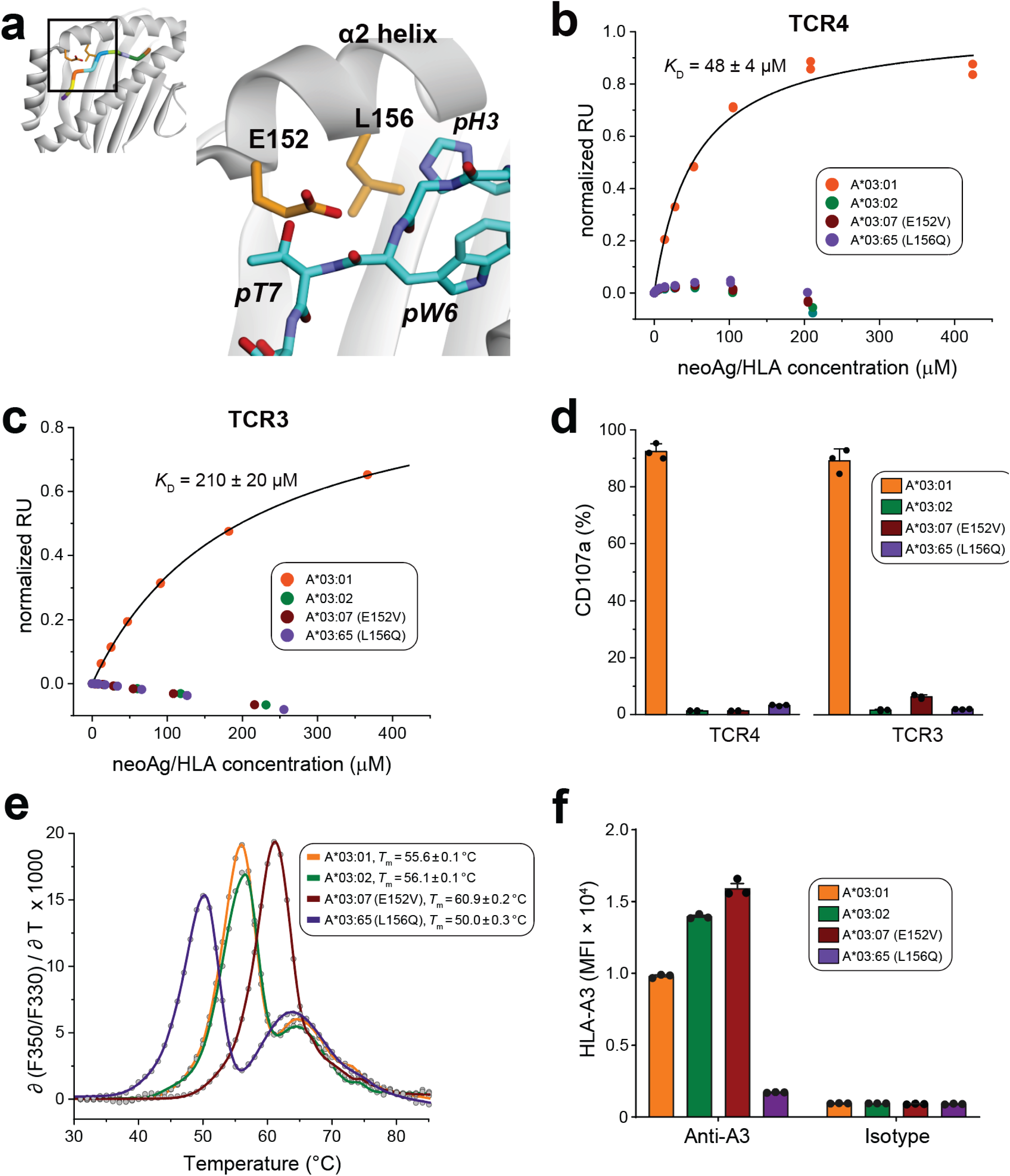
PI3Kα neoAg/HLA-A*03-specific TCRs distinguish between A*03:01 and A*03:02. **A)** Locations of positions 152 and 156 in the class I MHC protein. The two amino acids, Glu152 and Leu156 in A*03:01, are in the long arm of the α2 helix, as shown in the zoomed in region. **B)** Although binding to the neoAg/A*03:01 complex is readily detectable via SPR, the L156Q and E152V polymorphisms together or separately eliminate detectable binding of the neoAg specific receptor TCR4. The *K*_D_ value for TCR4 binding the A*03:01 complex is the average and standard deviation of three replicate measurements. **C)** As in panel B, but with TCR3. Together or separately, the two polymorphisms eliminate detectable binding. The *K*_D_ value for TCR3 binding the A*03:01 complex is the average and standard deviation of three replicate measurements. **D)** Together or separately, the polymorphisms also eliminate recognition of the neoAg in functional assays measuring the degranulation marker CD107a, although recognition when presented by A*03:01 is readily detectable. Data are absolute frequencies normalized for HLA expression using an allo-reactive control TCR ^90^; points and error bars are means and SEM from three replicates. **E)** The affinity of the neoAg for A*03:01 and A*03:02 is identical as measured by DSF. The E152V polymorphism alone improves peptide binding, whereas the L156Q polymorphism alone weakens it. *T*_m_ values are the average and standard deviation of 12 replicate measurements. **F)** Surface expression of individual HLA alleles matches stability of the recombinant neoAg/A3 complexes. Points and error bars are means and SEM from three replicates.

To assess TCR binding biochemically, we examined recognition using surface plasmon resonance (SPR), focusing on two neoAg-specific TCRs we previously characterized in detail. These TCRs, termed TCR3 and TCR4, recognize the neoAg/A*03:01 complex with high avidity and mediate the killing of A*03:01+ but not A*03:02+ targets presenting the PI3Kα neoAg ^33^. Consistent with the functional data, neither TCR showed detectable binding to the neoAg/A*03:02 complex by SPR, although binding to the A*03:01 complex was readily quantifiable (**Figs. 1B, C**). The neoAg/HLA*03:02 complex in this experiment was properly folded and assembled, confirmed by its binding to the positive control molecule S3-4, a single chain TCR variant (scTv) that binds peptide/HLA complexes on the “side” and largely independent of the bound peptide (**Fig. S1**) ^45^.

To explore the individual impacts of the two polymorphisms, we generated both A*03:07 (E152V) and A*03:65 (L156Q) and tested their recognition when presenting the neoAg. The binding of TCR3 and TCR4 to both complexes was again undetectable via SPR, despite both being recognized by the S3-4 scTv positive control (**Figs. 1B, C, and S1**). Neither complex was recognized in functional assays of degranulation measuring CD107a expression, mirroring the behavior of the A*03:02 complex (**Fig. 1D**). To assess the impact of the two polymorphisms on peptide binding affinity, we measured the thermal stabilities of the complexes using differential scanning fluorimetry (DSF), relying on the correlation between peptide binding and peptide/MHC thermal stability ^46,47^. As found previously, the thermal stabilities (*T*_m_ values) and thus peptide binding affinities of the A*03:01 and A*03:02 complexes were identical. The two polymorphisms had opposing effects, with E152V improving stability by ∼5 °C and L156Q weakening it by ∼5 °C, demonstrating an epistatic coupling between the polymorphic sites (**Fig. 1E**). To confirm this finding, we generated a set of transiently transfected cells expressing each individual A*03 allele and observed cell surface expression that strongly correlated with their measured thermal stability (**Fig. 1F**).

### Static structures fail to explain TCR selectivity between A*03:01 and A*03:02

To ask if there are structural differences between the neoAg complexes with A*03:01 and A*03:02 that could explain TCR selectivity, we crystallized and determined the structure of the neoAg/A*03:02 complex (**Table S1**). The structure is essentially identical to our previously determined structure of the neoAg/A*03:01 complex: the peptides superimpose with Cα and all atom root mean square deviations (RMSDs) of 0.3 and 0.7 Å, respectively (**Fig. 2A**). The peptide binding domains (residues 1-180) superimpose with Cα and all common atom RMSDs of 0.2 and 0.5 Å, respectively. There are no conformational perturbations at the sites of the two polymorphisms (E152V and L156Q), with the A*03:02 side chains overlaying those in A*03:01. The only apparent differences are in the interatomic interactions made via the two polymorphisms, as they alter a network of electrostatic interactions with the peptide and HLA protein (**Fig. 2B**). E152V results in the loss of a salt bridge with Arg114 in the base of the HLA-A3 binding groove. L156Q, on the other hand, introduces a new hydrogen bond with Arg114. Aside from these changes in electrostatic interactions, there are no structural changes in response to the two polymorphisms.

**Figure 2.**
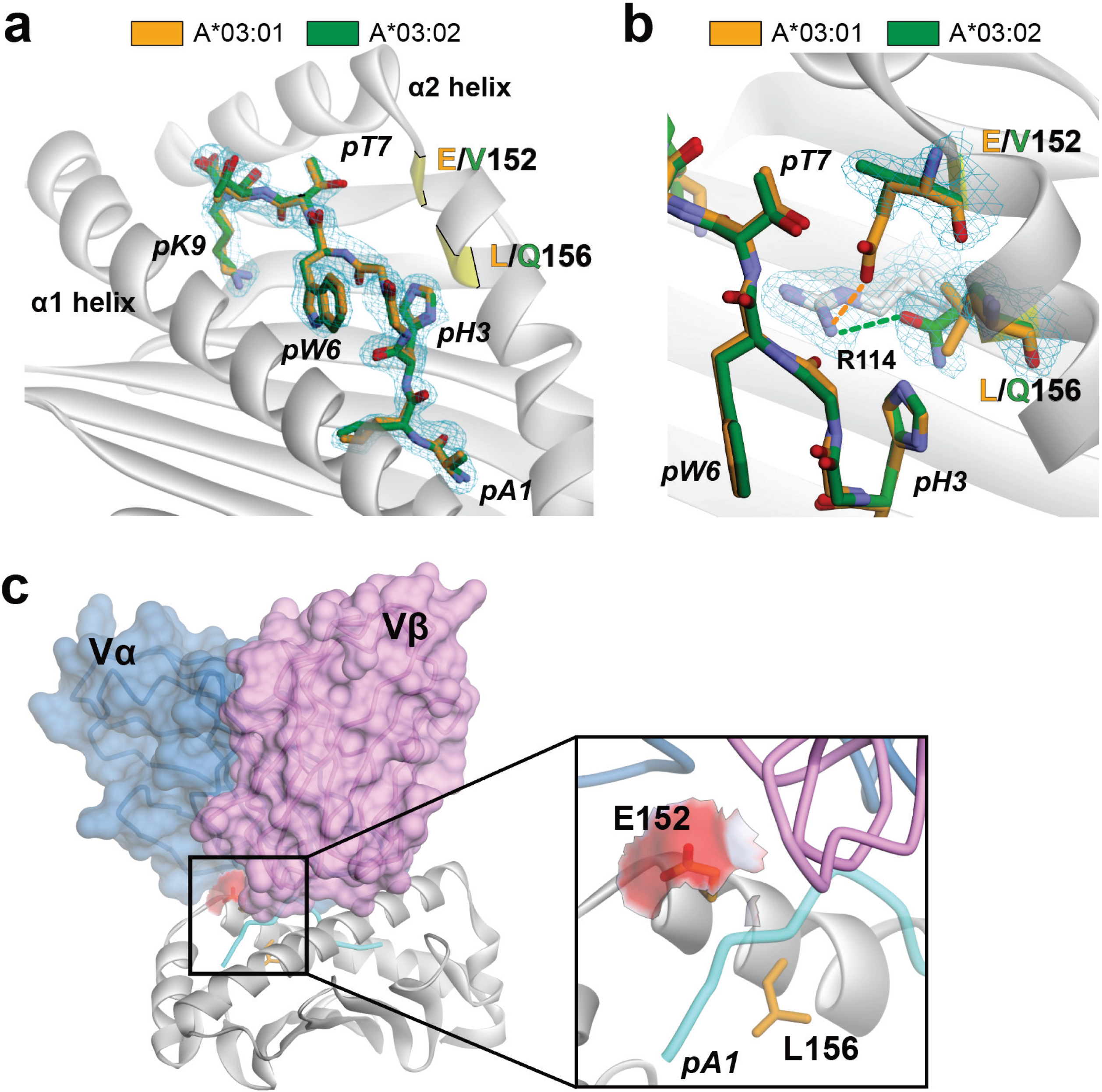
Static structures do not explain TCR selectivity between A*03:01 and A*03:02. **A)** The crystallographic structure of the PI3Kα neoAg/A*03:02 complex is identical to the structure of the neoAg/A*03:01 complex. The image shows the all atom superimposition of the peptides in the peptide binding groove. Cα and all atom RMSDs are 0.3 and 0.7 Å, respectively. The peptide binding domains superimpose with Cα and all common atom RMSDs of 0.2 and 0.5 Å, respectively. The mesh shows 2F_o_-F_c_ electron density for the peptide from a composite OMIT map calculated with simulated annealing, contoured at 1σ. The positions of the polymorphisms distinguishing A*03:02 from A*03:01 are indicated. **B)** As in panel A but zoomed in to the region of the polymorphisms. At position 152, the common atoms of the valine in A*03:02 overlay those of the glutamate in A*03:01, with the E152V polymorphism removing the salt-bridge with Arg114 in the base of the groove. At position 156, the new glutamine in A*03:02 overlays the leucine in A*03:01, with the L156Q polymorphism introducing a new hydrogen bond with Arg114. **C)** In the complex of TCR3 bound to the neoAg/A*03:01 complex (PDB 7RRG), Glu152 is fully exposed. The zoomed in panel shows the solvent accessible surface area of the Glu152 side chain, colored by atom identity (red = oxygen, white = carbon). There are no hydrogen bonds or longer-range electrostatic interactions made by the TCR to the exposed Glu152 side chain.

Given the absence of any conformational differences between the neoAg complexes with A*03:01 and A*03:02, we then asked if the polymorphisms could impact TCR recognition by influencing interactions in the TCR binding interface. We previously determined the structures of TCR3 and TCR4 bound to the neoAg/A*03:01 complex ^33^. Analyses of these structures showed minimal contact with the polymorphic residues. Neither TCR interacts with Leu156, which lies deep in the peptide-binding groove, away from the TCRs binding interface. While TCR4 interacts with Glu152 via a hydrogen bond from Thr96 in CDR3α, in the TCR3 complex, Glu152 is solvent exposed and does not form any short or long-range interactions with TCR3 (**Fig. 2C**). These findings indicate that static structures alone are insufficient to explain the TCRs selectivity between A*03:01 and A*03:02.

### NeoAg motional properties differ between A*03:01 and A*03:02

TCR binding to the PI3Kα neoAg in the context of A*03:01 requires a dramatic peptide conformational change, where the central tryptophan at position 6 (pTrp6) “flips” from lying along the α1 helix side of the binding groove to the α2 helix side (**Fig. S2**) ^33,34^. The flip occurs via a mechanism that sees the side chain rotating underneath and around the peptide backbone, moving through a large solvent filled cavity that exists between the peptide and the floor of the A*03:01 heavy chain. This conformational change, among the largest seen in TCR recognition of a peptide/MHC complex, is facilitated by high frequency peptide dynamics in the A*03:01 complex, with the tryptophan side chain sampling a range of conformations around its static crystallographic structure, including partially flipped conformations that position the side chain within the cavity near the base of the peptide binding groove ^34^.

The large cavity is also present in A*03:02, where it is occupied by multiple crystallographically resolved water molecules (**Fig. 3A**). Although the potential for a conformational flip in the peptide is thus present, we were curious if the E152V and L156Q polymorphisms might sterically hinder a flipped conformation. We superimposed the stretch of the α2 helix with the polymorphisms from the new neoAg/A*03:02 structure onto the same region of A*03:01 in the structure of the ternary complex with TCR3. There were a small number of steric clashes between the mutated side chains and the flipped peptide (**Fig. 3B**). Notably though, there are similar clashes when the same region from the TCR-free (apo) neoAg/A*03:01 structure is similarly superimposed into the ternary complex (**Fig. 3C**). With A*03:01, upon TCR binding these clashes are resolved by small side chain rearrangements and a shift in the α helix geometry. These are not unusual in TCR recognition of peptide/MHC complexes ^48^, and similar shifts would resolve the putative conflicts with A*03:02. However, the results of this modeling suggest that TCR discrimination between the neoAg presented by A*03:01 and A*03:02 could be attributable to polymorphism-dependent changes in the ability of the peptide to structurally move in a manner required for TCR binding.

**Figure 3.**
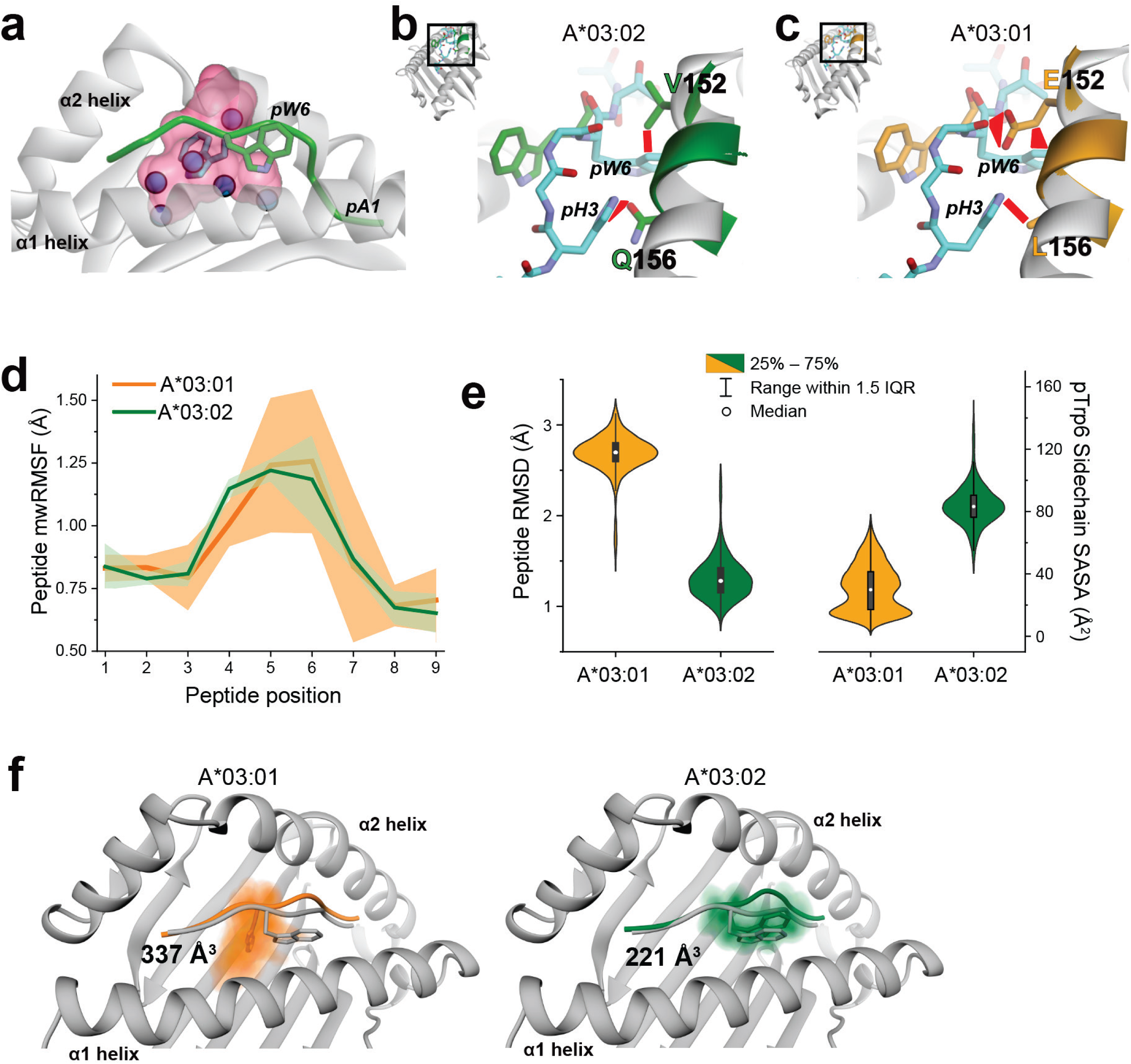
The PI3Kα neoAg samples a smaller conformational ensemble with an exposed pTrp6 when bound to A*03:02 compared to A*03:01. **A)** The large cavity between the neoAg backbone and the MHC binding groove seen in the structure with A*03:01 is also present in the structure with A*03:02. The cavity in the A*03:02 complex is shown in transparent pink. Crystallographically resolved water molecules within the cavity are shown as blue spheres. **B)** Modeling the α2 helix from the neoAg/HLA-A*03:02 complex (amino acids 152-157, Cα superimposition) into the ternary complex of TCR3 bound to the neoAg/A*03:01 complex indicates the potential for steric clashes (red polygons) with polymorphic sites if the TCR were to bind to the A*03:02 complex and the peptide were to flip. The sections from the apo neoAg/A*03:02 complex are in green (only pTrp6 is shown for the peptide). The conformations of A*03:01 are grey for the heavy chain and cyan for the peptide. **C)** As with panel B but superimposing the same stretch of the α2 helix from the apo neoAg/A*03:01 complex. Similar clashes are seen; with the A*03:01 ternary complex these are resolved with subtle side chain rearrangements and a small shift in helical geometry. **D)** Molecular dynamics simulations indicate the neoAg samples a smaller conformational ensemble when bound to A*03:02 compared to A*03:01. Solid lines are the average mass-weighted RMS fluctuations from the replicate 2 μs simulations (3 replicates for A*03:02; 4 for A*03:01); shaded areas give the standard deviations. The smaller variance for A*03:02 indicates more limited conformational sampling. **E)** Left panel: more limited sampling in A*03:02 is also apparent by examining RMS deviations from the starting configuration during the simulations: for A*03:02, the peptide remains close to its crystallographic conformation, whereas in A*03:01 it adopts divergent conformations. Right panel: in A*03:02, the more limited conformational sampling leaves the pTrp6 side chain more solvent exposed as shown by measurements of solvent accessible surface areas across the simulations. **F)** Volume occupied by pTrp6 of the neoAg during the simulations. Color density reflects degree of sampling; value gives volume of sampled space. Colored peptide is the average conformation; grey is the crystallographic conformation. In A*03:01 (left) pTrp6 samples conformations within the base of the binding groove. In A*03:02 (right) pTrp6 remains close to its solvent exposed crystallographic conformation. For panels D-F, simulation data for A*03:01 is from ref. 34.

We thus used molecular dynamics (MD) simulations to compare the motional properties of the neoAg in complex with A*03:01 and A*03:02. In our previous work, we examined the A*03:01 complex with lengthy, replicate atomistic simulations ^34^. We simulated the A*03:02 complex with the same protocol, performing three independent 2 μs simulations. From these, we first examined the root mean square fluctuations (RMSFs) of the peptides. The results indicated that, overall, the neoAg sampled fewer conformations when bound to A*03:02 compared to A*03:01: across the replicate simulations, the average peptide RMSFs were similar, but the variance was substantially lower for A*03:02 compared to A*03:01, particularly for positions 4-7 of the peptide (**Fig. 3D**). We then computed the root mean square deviations (RMSDs) of the peptides from their starting configurations. Across the replicate simulations, the values were much smaller for the A*03:02 complex, consistent with more limited conformational sampling (**Fig. 3E, left**). In our prior work with A*03:01, we observed that pTrp6 sampled conformations that buried the side chain in the base of the groove ^34^. We thus analyzed the solvent accessibility of the pTrp6 side chain during the simulations. In A*03:02 the side chain was substantially more solvent exposed, indicating a reduced tendency for the side chain to move into the base of the groove, reflecting, more restricted motion around the crystallographically observed conformation compared to A*03:01 (**Fig. 3E, right**).

To visualize the conformations sampled during the MD simulations, we defined 3D grids around pTrp6 with a spacing of 0.1 Å and tabulated the occupancy of each resulting voxel by the pTrp6 atoms throughout the simulations. The results showed that whereas pTrp6 in the A*03:01 complex sampled a large volume that extended into the base of the binding groove (**Fig. 3F, left**), in A*03:02 pTrp6 sampled a smaller volume and indeed remained close to the exposed conformation seen in the crystal structure (**Fig. 3F, right**).

### Experimental observation of altered peptide dynamics via ^19^F NMR

To experimentally validate the conclusions of the molecular dynamics simulations, we examined the motional properties of pTrp6 with nuclear magnetic resonance (NMR). We previously used NMR to study the motions of the PI3Kα neoAg bound to A*03:01, using a peptide variant in which pTrp6 was labeled with an NMR-active ^19^F atom ^34^. Consistent with the results of molecular dynamics simulations, those experiments showed that the pTrp6 side chain in A*03:01 samples a range of interconverting conformations that place the pTrp6 side chain in both buried and solvent-exposed positions. This was evident as multiple, broad peaks in the 1D NMR spectrum of the ^19^F-labeled neoAg/A*03:01 complex corresponding to alternative peptide conformations.

We performed these same experiments with the ^19^F-labeled neoAg/A*03:02 complex. Bound to A*03:02, the peptide showed only a single peak, indicating the side chain remains predominantly in one chemical environment and does not visit alternative states to a detectable extent (**Fig. 4**). The position of this lone peak in the A*03:02 spectrum was close to that of the major peak in the A*03:01 spectrum (−124.93 ppm in A*03:02, compared to −125.18 ppm in A*03:01), indicating similar chemical environments. Notably, the position of this peak was close to that of the free (unbound to HLA) ^19^F-labeled neoAg (−120.05 ppm), implying a solvent exposed environment. The NMR data thus indicate that, when bound to A*03:02, pTrp6 of the neoAg is conformationally restricted to a predominantly solvent exposed state compared to pTrp6 in the A*03:01 complex, fully consistent with the MD simulations.

**Figure 4.**
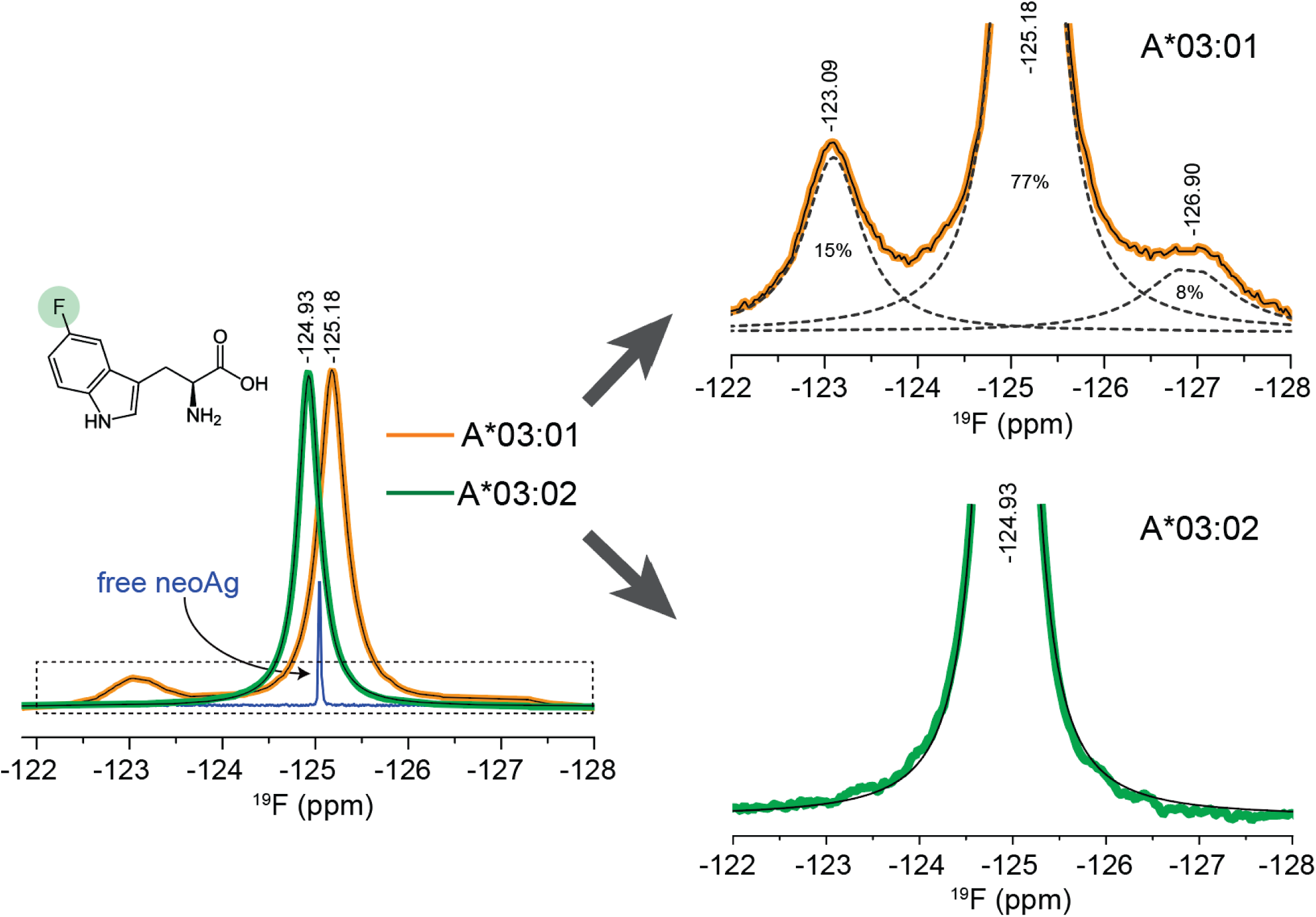
NMR confirms the differential mobility of the pTrp6 side chain in the context of A*03:02 compared to A*03:01. The figure shows 1D NMR spectra for the neoAg with an ^19^F-labeled Trp (inset) at position 6 bound to A*03:02 and A*03:01. The sharper blue line shows the data for the free neoAg in the absence of HLA, scaled vertically for clarity. The panels on the right show the indicated zoomed in regions for the two complexes. The more limited sampling in A*03:02 is clear from the reduced complexity of its spectrum compared to that of A*03:01. Notably missing from the A*03:02 spectrum is the peak near −123.09 ppm that we previously assigned to a buried pTrp6 conformation. Data for A*03:01 is from ref. 34.

### Altered motions resulting from changes to electrostatics block the peptide flip and TCR recognition of A*03:02

The data thus far indicate that the polymorphisms distinguishing A*03:02 from A*03:01 alter electrostatic interactions between the peptide, HLA α2 helix, and floor of the binding groove, resulting in altered motional properties of the PI3Kα neoAg. We next asked whether these changes impact the ability of the pTrp6 to flip upon TCR binding. To test this we used a steered molecular dynamics technique referred to as enforced rotation, which uses steering forces to move atoms around a rotational axis while permitting unbiased motions throughout the rest of the protein ^49^. We previously used this approach to assess how the identity of the position 2 anchor impacts the ability of the neoAg to flip when bound to A*03:01, applying force to pTrp6 to rotate it around the axis of the peptide backbone ^34^. We repeated those simulations here, this time examining the neoAg bound to A*03:02. With force constants ranging from 200 to 1600 kJ/mol/nm^2^ and attempting to rotate pTrp6 under the axis of the peptide backbone and through the cavity, we found that the neoAg bound to A*03:02 could not be forced to flip to the conformation required for TCR binding. In each case, the side chain quickly moved and remained 8-9 Å away from its flipped conformation, ending with the pTrp6 exposed and the peptide partially dissociating at the N-terminus (**Figs. 5A, B**). In contrast, the neoAg in the context of A*03:01 readily adopted the TCR-bound flipped conformation using even lower force constants (**Fig. 5A**) ^34^.

**Figure 5.**
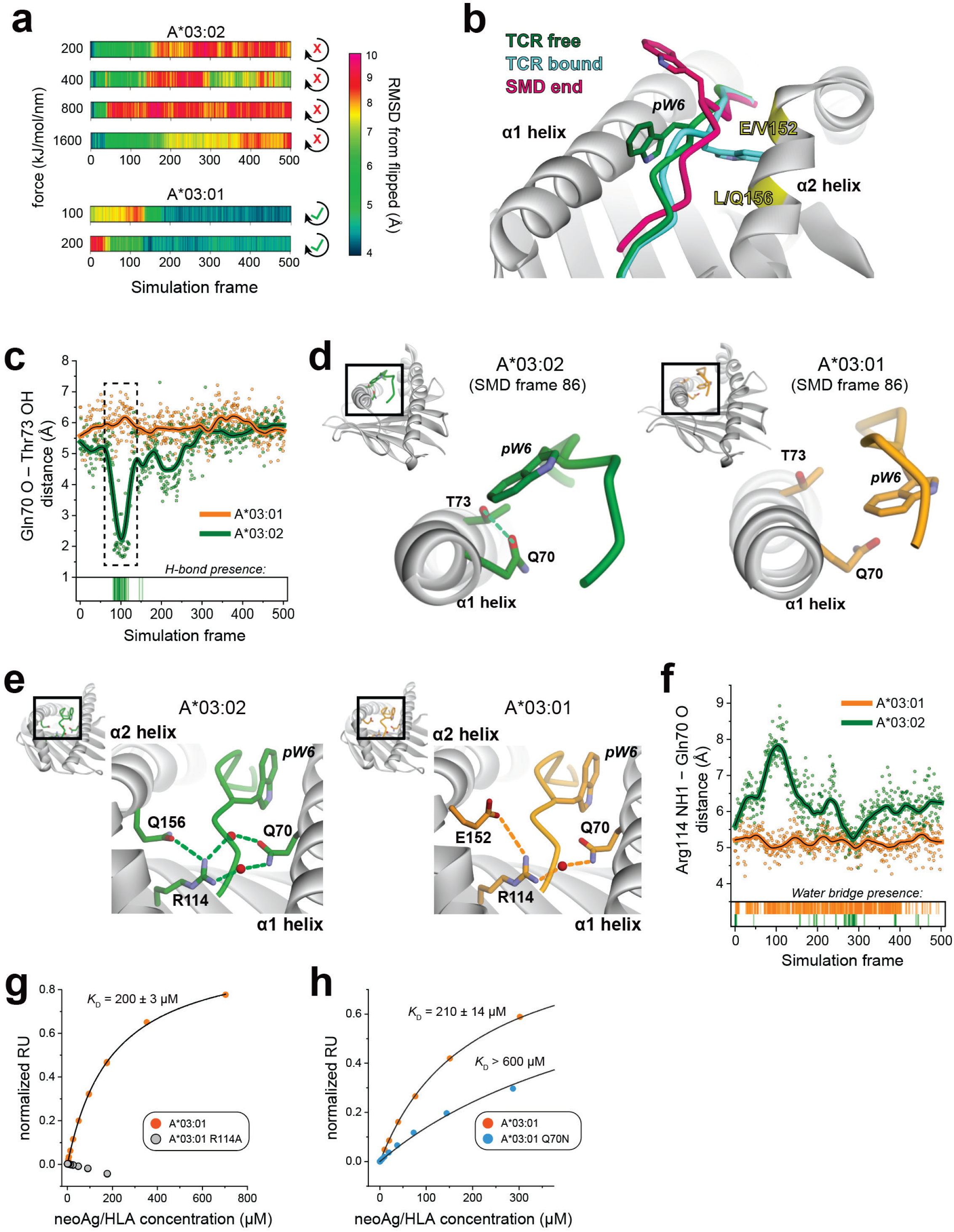
Altered motions resulting from changes to electrostatic interactions block the peptide flip and TCR recognition of A*03:02. **A)** Steered molecular dynamics is unable to rotate the neoAg in the context of A*03:02 into a TCR-bound flipped conformation using steering forces from 200 kJ/mol/nm^2^ to 1600 kJ/mol/nm^2^. In contrast, the neoAg in the context of A*03:01 readily rotated at even lower forces as shown at the bottom. Values show the pTrp6 RMSD from the flipped conformation in TCR4 according to the scale on the right. **B)** The final conformation of the neoAg in the A*03:02 steered molecular dynamics simulation with the 200 kJ/mol/nm^2^ force constant leaves pTrp6 exposed and the N-terminus partially dissociated. The final conformation is magenta, the starting conformation green, and the conformation bound to TCR4 cyan. **C)** In the SMD simulations with A*03:02, as force is applied, Gln70 rotates to hydrogen bond with neighboring Thr73, as indicated by the close distance between the Gln70 side chain oxygen and the hydroxyl group of Thr73 in the boxed region spanning frames 60-150. This rotation and hydrogen bond does not occur with A*03:01. The barcode at the bottom indicates the presence of the Gln70-Thr73 hydrogen bond in A*03:02. Data are from the simulations with the 200 kJ/mol/nm^2^ force constant. Solid lines are LOWESS smoothing of the datapoints. **D)** Movement of Gln70 in A*03:02 to hydrogen bond with Thr73 sterically impedes the movement of pTrp6 into the groove (left panel). This does not occur with A*03:01 and pTrp6 has begun its transition (right panel). Structural snapshots are from frame 86 of the 200 kJ/mol/nm^2^ simulations. **E)** Gln70 of the α1 helix forms water-bridged hydrogen bonds to Arg114 in both the A*03:02 and A*03:01 crystallographic structures with the neoAg. In A*03:02 (left panel), Arg114 in turn forms a hydrogen bond with Gln156. In A*03:01, Arg114 forms a salt bridge with Glu152 (right panel). **F)** Compared to A*03:01, the weaker connection to Arg114 in A*03:02 results in more frequent water dissociation and “untethering” of the connection with Gln70 in the SMD simulations, as shown by the increased distance between the Gln70 oxygen and Arg114 NH1, permitting formation of the Gln70 hydrogen bond with Thr73 as shown in panels C and D. The barcodes at the bottom indicate the presence of a water-bridged hydrogen between Arg114 and Gln70, with a frequency of 61% for A*03:01 and 8% for A*03:02. Data are from the simulations with the 200 kJ/mol/nm^2^ force constant. Solid lines are LOWESS smoothing of the datapoints. **G)** Introducing the R114A mutation into A*03:01 eliminates detectable binding of TCR3 to the neoAg/A*03:01 complex as shown by SPR. **H)** Introducing the Q70N mutation into A*03:01 weakens binding of TCR3 to the neoAg/A*03:01 complex by at least three-fold as shown by SPR. The *K*_D_ values for TCR3 binding are the averages and standard deviations of 2-5 replicate measurements.

The E152V and L156Q polymorphisms that alter pTrp6 motions and hinder its flip are in the α2 helix, on the opposing side of the peptide binding groove from where pTrp6 of the neoAg sits alongside the α1 helix in the TCR-free complex. In asking how the polymorphisms achieve their long-distance, cross-groove impact, we observed that in the steered MD simulations with A*03:02, as force is applied to pTrp6 to move it into the groove, Gln70 of the α1 helix moves up from its initial position, impeding the path pTrp6 takes to enter the base of the groove. This is facilitated by formation of a hydrogen bond with neighboring Thr73. Neither the Gln70 upward motion nor a hydrogen bond with Thr73 are observed in the simulations with A*03:01, and pTrp6 can enter and eventually transit the groove (**Figs. 5C, D**).

Asking how positions 152 and 156 in the α2 helix connect to Gln70 in the α1 helix, we noted that, in the crystal structures of the neoAg with both A*03:01 and A*03:02, Gln70 makes water-bridged hydrogen bonds with Arg114 in the base of the groove (**Fig. 5E**). Arg114 is in turn linked to positions 152 and 156: as noted earlier, in A*03:01, Arg114 forms a salt-bridge with Glu152. In A*03:02, the E152V polymorphism removes this salt-bridge, but it is replaced with a hydrogen bond when Leu156 is changed to glutamine (**Figs. 2B, 5E**). Thus, via buried water and Arg114, Gln70 is electrostatically connected to the sites of the polymorphisms, through a salt-bridge to Glu152 in A*03:01, versus a presumably weaker hydrogen bond to Gln156 in A*03:02.

To explore the consequences of this change in electrostatic interactions, we examined the water-mediated connectivity between Arg114 and Gln70 in the steered MD simulations. In the A*03:01 simulations, Arg114 remains tethered to Gln70 via one or more long-resident water molecules. These waters dissociate in the A*03:02 simulations, increasing the distance and reducing the tethering between Arg114 and Gln70 (8% tethering in the 200 kJ/mol/nm^2^ simulation with A*03:02, compared to 61% with A*03:01) (**Fig. 5F**). The resulting “untethering” from Arg114 allows Gln70 to sample alternative conformations, one of which includes an upward transition towards the pTrp6 side chain, impeding its rotation into the groove.

The data indicate a mechanism whereby the polymorphisms in the α2 helix that distinguish A*03:02 from A*03:01 block TCR recognition by altering the interactions with the key mediator Arg114, replacing a salt bridge with a weaker hydrogen bond. This in turn alters interactions in the groove, with one consequence being that, in A*03:02, a more weakly tethered Gln70 in the α1 helix moves to hydrogen bond with Thr73, sterically hindering the flip of the pTrp6 (as Arg114 also interacts directly with the peptide to facilitate its transition in the groove with A*03:01, other conformational consequences emerging from the weaker tethering to Arg114 are likely).

To test this mechanism experimentally, we replaced Arg114 with alanine in the context of A*03:01 (i.e., retaining glutamate at position 152 and leucine at position 156). We were unable to detect binding of TCR3 via SPR (**Fig. 5G**), although the R114A mutant complex was well recognized by the peptide-independent S3-4 scTv positive control and thus properly assembled (**Fig. S1**). For a more subtle perturbation, we changed Gln70 on the α1 helix to asparagine in the context of A*03:01. This weakened binding TCR by at least 3-fold, consistent with a weakening of the cross-groove connection (**Figs. 5H, S1**).

### The single polymorphisms that define A*03:07 and A*03:65 similarly impede TCR binding

We revisited the E152V and L156Q polymorphisms individually, which define the low-frequency alleles A*03:07 and A*03:65. As shown above, the PI3Kα neoAg is not biochemically or functionally recognized in the context of either of these proteins (**Figs 1A, C**). To ask if there was a shared, underlying mechanism with A*03:02 in which pTrp6 of the peptide could not flip to support TCR binding, we determined the crystallographic structures of the neoAg with both A*03:07 and A*03:65 (**Table S1**). The A*03:07 complex was essentially identical to the A*03:01 and A*03:02 complexes, with the E152V polymorphism removing the salt-bridge to Arg114 without structural consequences (**Fig. 6A**; peptide Cα and all atom RMSDs to the A*03:02 structure of 0.1 Å and 0.4 Å, respectively).

**Figure 6.**
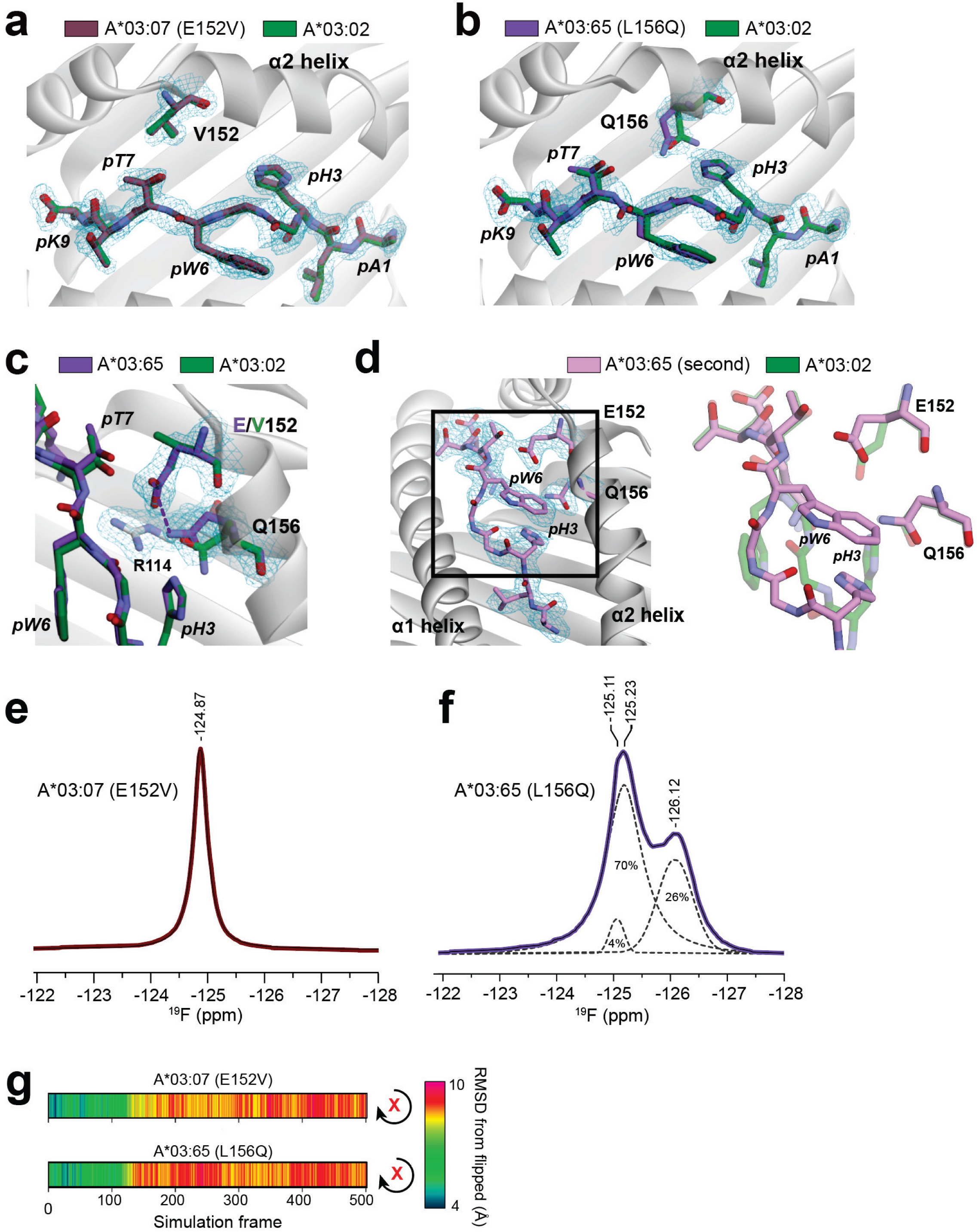
The polymorphisms that define A*03:07 and A*03:65 also impede TCR binding. **A)** Other than replacement of the side chain, the structure of the neoAg/A*03:07 (E152V) complex is identical to the structure of the neoAg/A*03:02 complex. The image shows the all atom superimposition of the peptides in the peptide binding groove. Cα and all atom peptide RMSDs are 0.1 and 0.4 Å, respectively. **B, C)** The structure of the first form of the neoAg/A*03:65 (L156Q) complex is also similar to the neoAg/A*03:02 complex, except that Gln156 is more recessed in the groove. This removes the hydrogen bond with Arg114 and replaces it with a hydrogen bond to Glu152, which no longer salt bridges with Arg114. Cα and all atom peptide RMSDs are 0.3 and 0.6 Å, respectively. **D)** The structure of the second form of the neoAg/A*03:65 (L156Q) complex reveals a substantially altered conformation of the peptide with an altered backbone and side chain placement for pTrp6. The position of Glu152 is also altered and Gln156 is recessed, eliminating the interactions between them and with Arg114. Cα and all atom peptide RMSDs are 1.5 and 2.8 Å to the first form and 1.4 and 2.5 Å to the structure with TCR3 bound. **E)** The ^19^F NMR spectra of the ^19^F-labeled neoAg/A*03:07 (E152V) complex is identical to that of the A*03:02 complex, indicative of very similar conformational behavior for pTrp6. **F)** Consistent with the crystal structures, the ^19^F NMR spectra of the ^19^F-labled neoAg/A*03:65 (L156Q) complex was more complex but lacked the downshifted peak indicative of the pTrp6 side chain buried in the groove. **G)** Steered molecular dynamics is unable to rotate the pTrp6 into a flipped conformation with A*03:07 or A*03:65, consistent with the cross-groove perturbations seen crystallographically. Data is for a 200 kJ/mol/nm^2^ force constant. For panels A-D, meshes show 2F_o_-F_c_ electron densities from composite OMIT maps calculated with simulated annealing, contoured at 1σ.

Unlike the other complexes, the A*03:65 complex crystallized in two different forms, the first form nearly identical to that of the others and the second form with a different precipitant and buffering reagent. We determined the structures from both crystal forms (**Table S1**). The first was again similar to the A*03:01 and A*03:02 complexes (peptide Cα and all atom RMSDs to the A*03:02 structure of 0.3 Å and 0.6 Å, respectively), but compared to A*03:02, the L156Q polymorphism led to Gln156 lying more recessed in the groove, removing the hydrogen bond with Arg114 and replacing it with a hydrogen bond to Glu152. This leads to a shift that also removes the salt bridge with Arg114 (**Figs. 6B, C**). The structure of the second form was different, showing a substantially altered peptide conformation, with the pTrp6 pointed towards the α2 helix and the peptide backbone occupying the space formerly filled by the pTrp6 side chain (peptide Cα and all atom RMSDs to the first crystal form of 1.5 Å and 2.8 Å). The peptide conformation in this altered form is also very different from that in the flipped, TCR-bound conformation (Cα and all atom RMSDs to the form in complex with TCR3 of 1.4 Å and 2.5 Å, respectively). In this second form, the position of Glu152 is also altered and Gln156 remains recessed as seen in the first form. Together, these eliminate the interactions between them and with Arg114 (**Fig. 6D**).

Although the two single mutations differentially affect the structure and stability of the complexes, common to both is a loss of interactions with Arg114 at the base of the peptide-binding groove, the key mediator of the cross-groove mechanism that permits a peptide flip with A*03:01. To assess the effect of this change on peptide dynamics in the binding groove, we examined the A*03:07 and A*03:65 complexes with ^19^F-neoAg using ^19^F NMR as above. The 1D ^19^F NMR spectrum of the A*03:07 sample was almost identical to that of the A*03:02, showing a single peak at nearly the same position (−124.87 ppm in A*03:07 versus −124.93 ppm in A*03:02; **Fig. 6E**). This indicates that, as in A*03:02, when the neoAg is bound to A*03:07, the pTrp6 side chain remains predominantly solvent-exposed and conformationally restricted. The spectrum of the A*03:65 sample was more complex, revealing two major peaks (**Fig. 6F**). This is consistent with the crystallographic data showing at least two stable conformations available to the peptide when bound to A*03:65. While there was some overlap with the A*03:01 spectrum, the spectrum for A*03:65 notably lacked the peak near −123 ppm that we previously associated with the buried conformations of the pTrp6 side chain at the base of the peptide binding groove ^34^.

We next performed steered molecular dynamics simulations with the A*03:07 and A*03:65 complexes (using the structure from the first crystal form for A*03:65) and were again unable to induce peptide rotation (**Fig. 6G**). As in the simulation with A*03:02, the water-bridged connection between Arg114 and Gln70 is lost early in these simulations as force is applied (**Fig. S3**). We conclude that, while fine details may differ, the general mechanism by which A*03:07 and A*03:65 eliminate recognition of the PI3Kα neoAg is the same as in A*03:02; i.e., an inability for the pTrp6 to transit under the peptide backbone and adopt a TCR-binding competent conformation due to the perturbation of a cross-groove electrostatic linkage that relies on Arg114.

### The cross-groove electrostatic linkage is present in other structures with A*03:01 but absent in other members of the HLA-A3 superfamily

Lastly, we examined other high resolution peptide/HLA structures with resolved waters to ask about the prevalence of this cross-groove linkage. In other structures with A*03:01, we found the same water-bridged linkage between Gln70 in the α1 helix, Arg114 in the base of the groove, and Glu152 in the α2 helix in structures with epitopes from influenza, HIV, and the KRAS oncogene (PDB IDs 3RL1, 7MLE, 7UC5, and 8VJZ) ^20,50,51^. There are no structures available for TCRs bound to any of these peptide/A*03:01 complexes, but the peptide conformations are all extensively bulged with cavities and buried water between the peptide and the base of the binding groove. There are no other structures available with A*03:02, but a linkage resembling the perturbed A*03:02 linkage is in seen in multiple structures with HLA-A*11:01 (PDB IDs 1QVO, 4MJ5, 7M8T, 7OW4, 7S8Q, and 8RHQ) ^52–57^, a member of the HLA-A3 superfamily that retains Gln70 and Arg114 but replaces positions 152 and 156 with an alanine and a glutamine, respectively. In other members of the HLA-A3 superfamily, the linkage is missing altogether, as the various amino acids at positions 70, 114, 152, 156 are substituted by a wide variety of other amino acids.

## Discussion

The micropolymorphisms that distinguish closely related HLA proteins are associated with diverse outcomes, including differential risks of autoimmunity, hypersensitivity, control of viral infection, and transplant success. In addition to changes in peptide binding affinities and thus peptide repertoires, micropolymorphisms have been shown to alter static peptide/MHC conformation, TCR contact sites, and peptide or MHC molecular motions, all of which can influence TCR binding, specificity, and cross-reactivity. However, micropolymorphisms are often not considered in the context of the broader structural and dynamic properties of the peptide/MHC complex. Here we uncovered a novel mechanism through which micropolymorphisms can impact TCR selectivity, with the amino acid differences at positions 152 and 156 that distinguish A*03:02 from A*03:01 disrupting a cross-groove, interconnected network of polymorphic sites that allows the peptide/MHC complex to structurally adapt for optimal TCR binding. More complex than previously reported changes in peptide or MHC flexibility, this mechanism involves not only the peptide, but the flanking α helices, the base of the peptide binding groove, and coordinated interactions between them that facilitate conformational changes needed for TCR binding. Disruption of this cross-groove network turns a potent neoAg in the context of A*03:01 into a null antigen in the context of A*03:02, as well as in the context of the less frequent, singly substituted alleles A*03:07 and A*03:65.

Although our work examined differences between A*03:01 and A*03:02 and focused on the micropolymorphisms at positions 152 and 156 in the α2 helix, the other positions in the cross-groove network, position 114 in the base of the groove and position 70 in the α1 helix, all vary within and across class I superfamilies ^58,59^. Observations highlighting a strong influence of these positions on class I MHC functions date back decades ^60–62^. Interestingly though, across alleles, positions 70, 114, 152, and 156 display co-variance and are thus evolutionarily coupled ^63^. While one reason for co-variance likely involves their cooperative roles in defining the static protein architecture that influences what peptides bind and how they are displayed ^13–22^, our data suggest that this and other networks of amino acids are also co-selected to facilitate allele-dependent differences in protein dynamics and structural malleability, providing an additional means through which polymorphisms influence immunity. Further evidence for this is found in the influence of groups of coupled amino acids (including positions 114 and 156) on tapasin dependence, which requires significant protein conformational adaptability ^28,29,32,64–68^. Prior studies have also highlighted the presence of interconnected class I MHC polymorphic sites in other functions, including a connection between positions in the α1 and α2 helices in HLA-B*44 alleles that is also mediated by interactions with the base of the peptide binding groove ^15^. Overall, the evidence suggests that the class I MHC protein has evolved as an interconnected, dynamic system whose motions, adaptability, and thus immunological functions are tuned by interacting, co-varying networks of polymorphisms that helps maximize diversity in antigen presentation and recognition. In the specific alleles studied here, the role of evolutionary forces in selecting for conformational adaptability may be reflected in the greater frequency of A*03:01 over A*03:02 and other, less frequent members of the HLA-A3 superfamily such as A*03:07 and A*03:65.

Notably, with the PI3Kα neoAg, the micropolymorphisms in A*03:02 disrupt the cross-groove network without influencing either peptide binding affinity or the static structure of the unbound (apo) peptide/HLA complex. We believe this outcome is unlikely to be unique to the PI3Kα neoAg or members of the HLA-A3 superfamily. Observations of TCR specificity towards subtly different peptide/MHC complexes whose static structures are indistinguishable date back to some of the earliest work in structural immunology ^35–37^. The intersection of these findings with the frequent observations of peptide/MHC conformational changes upon TCR binding and the demonstrated high sensitivity of peptide/MHC dynamics to subtle changes suggests that the ability for recognition and thus antigenicity to be tuned via “structurally silent” mechanisms may not be unusual ^69–71^. This reflects the phenomenon of dynamic allostery – binding selectivity occurring through dynamic changes that often occur without discernable changes in structurally resolved “ground states” – that is becoming widely appreciated throughout biology ^72–74^. A clear example of this phenomenon with class I MHC proteins is illustrated by the murine Ly49C NK receptor that senses different peptides despite binding far from the peptide binding groove ^75^.

Our findings also bear on the role of applied forces in T cell signaling, particularly catch bonds that can form between TCRs and their targets in T cell – APC interfaces ^76–79^. While an influence of catch bonds is not implicated in our findings, the conclusion that micropolymorphisms tune the flexibility and adaptability of MHC-I proteins has implications for T cell mechanobiology. Although the fine details of individual catch bonds are poorly understood, the phenomenon in general depends on how the interacting proteins move and structurally evolve under force ^80,81^. The impact of class I MHC micropolymorphisms on structural adaptability is thus likely to introduce another complexity into how applied force and catch bonding influences immunogenicity.

In conclusion, our work has shown how the micropolymorphisms that distinguish A*03:01 from A*03:02 (and individually from A*03:07 and A*03:65) perturb an interconnected, cross-groove network of amino acids that, in A*03:01, facilitates a large conformational change required for TCR recognition of the PI3Kα neoAg. The perturbation leads to altered motions that hinder the conformational change, translating into allele specificity in neoAg recognition. Notably, this specificity is achieved without changes in peptide affinity or, for A*03:02 and A*03:07, without changes in the static structures of the unbound peptide/MHC complexes. Although focused on a high profile public neoAg from the PI3Kα oncoprotein, our results have implications across immunology, showcasing the flexibility and malleability of class I peptide/MHC complexes and how these have been evolutionarily tuned to promote diversity in antigen presentation and recognition, as well as amplify the physical impact of a small peptide in the context of the much larger MHC protein. Lastly, our findings reiterate calls for the use of high resolution HLA typing across T cell immunology, including in the rapidly evolving field of antigen-targeted immunotherapy.

## Acknowledgements

This research was supported by National Institutes of Health (NIH) awards R01AI17665 (to BMB), R35GM118166 (to BMB), and R37CA259177 (to CAK). CAK was also supported in part by R01CA286507, P50CA217694, P30CA008748, The Parker Institute for Cancer Immunotherapy, Cycle for Survival, and the Metropoulos Family Foundation. CAB was supported by NIH fellowship F32AI191525. BE was supported by the Notre Dame CBBI program, supported in part by NIH award T32GM145773. GP was supported in part by a fellowship from the Notre Dame Berthiaume Institute for Precision Health. The use of SPR was supported in part by NIH grant S10OD028553. This study also made use of the National Magnetic Resonance Facility at Madison, where NMR equipment, helium recovery equipment, and computers were purchased with funds from the University of Wisconsin-Madison and NIH awards P41GM136463, R24GM141526, P41GM103399, S10RR023438, S10RR025062, and S10RR029220. This work also used the GM/CA and NE-CAT beamlines at the APS. GM/CA@APS has been funded by the National Cancer Institute (ACB-12002) and the National Institute of General Medical Sciences (AGM-12006, P30GM138396). The NE-CAT beamlines have been funded by the National Institute of General Medical Sciences from the National Institutes of Health (P30GM124165). This research used a beamtime award (https://doi.org/10.46936/APS-190953/60014736) from the Advanced Photon Source, a U.S. Department of Energy (DOE) Office of Science User Facility operated for the DOE Office of Science by Argonne National Laboratory under Contract No. DE-AC02-06CH11357. We are grateful to Jean Custodio for structural biology assistance, Eliza Kovrigin for technical assistance, Dr. Marco Tonelli for providing pulse programs for NMR experiments and assisting with access to the NMR spectrometer, and the CCP4/APS School in Macromolecular Crystallography for training.

## Contributions

JM: structural biology; binding, stability, and NMR; traditional MD simulations; CMA: traditional MD simulations; CAB: SMD simulations; BE: binding and stability experiments; GP: structural biology; JAL: structural biology; ELK: NMR; SSC: T cell functional experiments; CAK: study design, funding; BMB: study design, funding; All authors: initial, intermediate, and final drafts.

## Methods

### Recombinant protein production

All peptides were purchased from GenScript at >80% purity and dissolved in DMSO prior to refolding. Recombinant plasmids with genes encoding the extracellular domain of MHC heavy chains, β_2_-microglobulin, TCR α and β chains, and the S3-4 scTv were either obtained commercially from GENEWIZ or generated by PCR-based site-directed mutagenesis. Proteins, including the MHC heavy chains, β_2_-microglobulin, TCR α and β chains, and the S3-4 scTv were then purified from bacterially expressed inclusion bodies. Briefly, the proteins were overexpressed in *Escherichia coli* and the resulting inclusion bodies solubilized in 8 M urea and 6 M guanidinium-HCl. Denatured proteins were refolded in either peptide/MHC refolding buffer (400 mM L-arginine, 100 mM Tris-HCl, 2 mM Na_2_EDTA, 6.3 mM cysteamine, 3.7 mM cystamine and 0.2 mM PMSF, pH 8.3) or TCR refolding buffer (50 mM Tris-HCl, 2.5 M urea, 2 mM Na_2_EDTA, 6.5 mM cysteamine, 3.7 mM cystamine and 0.2 mM PMSF, pH 8.15) at 4 °C and incubated overnight. The refolding buffer was then dialyzed against ddH_2_O followed by 10 mM Tris-HCl (pH 8.3) at room temperature (for peptide/MHC) or 4 °C (for TCR and scTv) for 48 h. The refolded proteins were subsequently purified by anion exchange followed by size exclusion chromatography. Protein concentrations were determined by UV absorbance at 280 nm using sequence-determined extinction coefficients.

### X-ray crystallography and structural analyses

Peptide/MHC complexes were solubilized in 10 mM HEPES, 20 mM NaCl, pH 7.4 prior to crystallization. Crystals were obtained by hanging drop vapor diffusion at 4 °C. Crystals of the neoAg/HLA-A*03:02 complex grew from 20% w/v polyethylene glycol 3350 and 200 mM lithium nitrate at pH 7.1. Crystals of the neoAg/HLA-A*03:07 (E152V) complex grew from 12% w/v polyethylene glycol 3350 and 100 mM sodium malonate at pH 6.0. For the neoAg/HLA-A*03:65 (L156Q) complexes, crystals of the first form grew from 20% w/v polyethylene glycol 3350 and 200 mM sodium malonate at pH 7.0, while the second form grew from 10% w/v polyethylene glycol 20000 and 100mM MES at pH 6.5. Crystals were cryoprotected with 15–25% glycerol prior to flash-freezing in liquid nitrogen. X-ray diffraction data were collected at the 23-ID-D (for the first form of neoAg/HLA-A*03:65 (L156Q) and 24-ID-C (for the other three structures) beamline of the Advanced Photon Source at Argonne National Laboratory. Diffraction data were processed through HKL2000 ^82^ and initially phased by molecular replacement using Phaser in Phenix ^83^. Space group determinations were confirmed by processing the data through DIALS followed by merging and scaling with Aimless in CCP4 ^84^. The search model for all structures was our previously solved neoAg/HLA-A*03:01 structure (PDB 7L1C) with the peptide removed. Peptides were then manually rebuilt in Coot after initial models were obtained from Phenix AutoBuild. Models were further refined automatically in Phenix and manually in Coot ^85^. Composite/iterative build OMIT maps were calculated with simulated annealing using CNS as implemented in Discovery Studio 2025. Structures were visualized and analyzed in PyMOL, Discovery Studio, VMD, and UCSF Chimera ^86,87^. Solvent accessible surfaces and surface areas were calculated in either Discovery Studio or VMD with a 1.4 Å probe radius. The cavity in the peptide/MHC complex was quantified with CAVER Analyst 2.0 with default options, recording the summed volumes of the largest contiguous pockets between the peptide and the MHC binding groove ^88^.

### SPR binding measurements

Binding affinities were measured via SPR using a Biacore T200 instrument. Proteins were buffer exchanged into HBS-EP buffer (10 mM HEPES, 150 mM NaCl, 3 mM EDTA, 0.005% surfactant P20, pH 7.4) prior to experiments. TCRs and the S3-4 scTv were immobilized on a CM5 Series S sensor chip to 1000-6000 RU via amine coupling and peptide/MHC complexes were injected at a flow rate of 5 μL/min as analytes. Experiments were performed at 25 °C with a blank activated and deactivated flow cell as reference. Binding affinities were determined by fitting the curves of the reference-subtracted steady-state responses against the injected protein concentrations to a 1:1 binding model in OriginPro 2024. In most cases injections at each concentration were repeated twice, and values from both sets of injections were fit simultaneously for a single measurement. With the Q70N mutation, global fitting with a shared surface activity was used to improve accuracy ^89^.

### T cell functional analysis

T cells were retrovirally transduced with either TCR3 or TCR4 as previously described ^34^. An alloreactive HLA-A3-binding TCR was included as a control to normalize for HLA expression ^90^. Neoantigen-specific activation of TCR- T cells was assessed by measuring upregulation of CD107a by flow cytometry. Cos-7 cells were electroporated with 100 μg/mL of *HLA-A*03:01, HLA-A*03:02, HLA-A*03:07, or HLA-A*03:65* mRNA and plated into 96-well round-bottom plates overnight at 37 °C to allow surface HLA expression. The following day, cells were pulsed with neoAg (1μg/mL) for 30 min at 37 °C. Cells were washed with 1× PBS to remove any unbound peptide. TCR-T cells were added at an E:T ratio of 1:1 for 6 h in the presence of anti-CD107A-BV650 (Clone H4A3, BioLegend). Cells were washed in 1× PBS and surface labeled with Live/Dead fixable dye (Invitrogen), anti-CD3-APC-H7 (Clone SK7, Invitrogen), anti-CD8-eFluor450 (Clone SK1, Invitrogen) and anti-mouse TCR-PerCpCy5.5 (Clone H57-597, Invitrogen) for 30 min at 4 °C. All antibodies were used at a final concentration of 5 μg/mL. Finally, cells were washed with 1× PBS, suspended in 2% FBS in PBS, and data acquired on an X20 LSR Fortessa flow cytometer with the BD FACSDiva software. Data were analyzed using FlowJo version 10.6.2.

### HLA surface expression

COS-7 cells were electroporated with 100 μg/mL of *HLA-A*03:01, HLA-A*03:02, HLA-A*03:07, or HLA-A*03:65* mRNA and plated into 96-well round-bottom plates overnight at 37 °C to allow surface HLA expression. The following day, cells were labeled with anti-HLA*A3 (Clone GAP.A3, Invitrogen) for 30 min at 4 °C. Cells were washed with 1x PBS, suspended in 2% FBS in PBS, and data acquired on an X20 LSR Fortessa flow cytometer with the BD FACSDiva software. Data were analyzed using FlowJo version 10.6.2.

### Differential scanning fluorimetry

Thermal denaturation of peptide/MHC complexes was performed using differential scanning fluorimetry using a Prometheus NT.48 instrument (NanoTemper) monitoring intrinsic tryptophan fluorescence ^47^. Briefly, 10 μL of each peptide/MHC complex at concentrations of 15-20 μM in HBS-EP buffer (10 mM HEPES, 150 mM NaCl, 3 mM EDTA, 0.005% surfactant P20, pH 7.4) were loaded into instrument capillaries. The temperature was scanned from 20 to 95 °C at a rate of 1 °C/min. Fluorescence at emission wavelengths of 330 nm and 350 nm was recorded, and the first derivative of the ratio of the fluorescence intensities was plotted against the temperature to generate melting curves. Data were fit to bi-Gaussian functions using OriginPro 2024.

### Molecular dynamics simulations

Fully atomistic molecular dynamics simulations of the neoAg/HLA-A*03:02 complex were performed on GPU hardware with Amber18 using the ff14SB force field and an SPC/E water model as previously described, except using the neoAg/A*03:02 complex ^33^. After minimization, production trajectories were calculated under constant volume and temperature with a 2 fs time step for a total time of 2 μs for each production trajectory. Trajectories were analyzed with CPPTRAJ 18 ^91^. Three independent replicate MD simulations were performed. The neoAg/HLA-A*03:01 complex was previously simulated using the same protocol in quadruplicate ^33^. Mass weighted amino acid RMSF values were calculated via the CPPTRAJ ‘atomicfluct’ command. 1D RMSD values were calculated via the CPPTRAJ ‘rms’ command after superimposition of the Cα atoms of the MHC binding groove (residues 1-180). Grid space occupancies were calculated via combinatorial usage of the CPPTRAJ ‘bounds’ and ‘grid’ commands using a grid spacing of 0.1 Å. Occupied grid space was visualized through the volume viewer in UCSF Chimera. The grid space volume was first smoothed via a Gaussian filter and contoured to envelope grid space occupied for approximately 10% of simulation time. Grid space calculations were performed using the first set of MD simulation data. Steered molecular dynamics (SMD) was performed using enforced rotation ^44^ as previously described, except using the structures of the neoAg/A*03:02, neoAg/A*03:07, and neoAg/A*03:65 complexes (the initial structure for A*03:65) ^33^. The rotation group was defined as all atoms of the pTrp6 amino acid and rotation vectors selected along the peptide backbone. Under-peptide enforced rotation was carried out for a set of force constants using the flex2-t potential and rotation rate of 0.1°/ps for 500 ps. Force constants used were 200, 400, 800, and 1600 kJ/mol/nm^2^. RMSD measurements of rotated pTrp6 coordinates relative to those observed in the TCR4-bound conformation (from PDB 7L1D) were obtained using the RMSD visualizer VMD plugin following a superposition of the Cα atoms of the residues 1-180 of the peptide binding grooves and selection of all pTrp6 atoms.

### Nuclear magnetic resonance

^19^F NMR spectra were recorded using a Bruker 600 MHz NMR instrument with a QCI cryogenic probe at the NMRFAM of the University of Wisconsin Madison as described previously ^34^. Data were collected at a calibrated value of 5.2 °C. Complexes were solubilized in 10 mM HEPES, 150 mM NaCl, pH 7.4 with 10% D_2_O (Cambridge Isotope Laboratories) at concentrations ranging from 0.3 mM to 0.5 mM. Pulse sequences, data collection and analysis were identical to the previous studies on A*03:01. The total acquisition times were 24 hours for A*03:02 and A*03:07 samples and 40 hours for the A*03:65 sample.

## Supplemental Figure Legends

**Figure S1.** SPR titrations of the positive control S3-4 scTv binding to various peptide/HLA-A*03 constructs. As indicated by the structural image at the top, S3-4 binds to the side of the HLA protein in a mostly peptide-independent fashion. Strong binding in each case demonstrates proper assembly of the peptide/HLA-A3 complex. *K*_D_ values are the average and standard error of at least two replicates.

**Figure S2.** In the context of A*03:01, the tryptophan at position 6 of the neoAg undergoes a dramatic flip upon TCR recognition. The image on the left illustrates the flip, with the TCR-bound conformation taken from the structure of TCR4 bound to the neoAg/HLA-A*03:01 complex. As shown on the right, the flip occurs via a mechanism whereby the pTrp6 side chain rotates underneath the peptide backbone. Figures adapted from ref. 34.

**Figure S3.** In the SMD simulations with A*03:07 and A*03:65, as seen with A*03:02, Gln70 rotates to hydrogen bond with Thr73 early in the simulation (boxed region), sterically hindering the pTrp6 flip.

**Table S1.** Crystallization and X-ray refinement statistics. Numbers in parentheses are for the highest resolution shell.

## References

1. Klebanoff, C.A., Chandran, S.S., Baker, B.M., Quezada, S.A. & Ribas, A. T cell receptor therapeutics: immunological targeting of the intracellular cancer proteome. Nature Reviews Drug Discovery (2023).

2. Doherty, P.C. & Zinkernagel, R.M. H-2 compatibility is required for T-cell-mediated lysis of target cells infected with lymphocytic choriomeningitis virus. J Exp Med 141, 502–7 (1975).

3. Robinson, J., et al. The IMGT/HLA database. Nucleic acids research 39, D1171–6 (2011).

4. Sidney, J., Peters, B., Frahm, N., Brander, C. & Sette, A. HLA class I supertypes: a revised and updated classification. BMC Immunology 9, 1 (2008).

5. Reveille, J.D. et al. HLA class I and II alleles in susceptibility to ankylosing spondylitis. Annals of the Rheumatic Diseases 78, 66–73 (2019).

6. Leccese, P., et al. The relationship between HLA-B*51 subtypes, clinical manifestations and severity of Behçet’s syndrome: a large Italian cohort study. Rheumatol Adv Pract 7, rkad087 (2023).

7. Mikk, M.L. et al. The HLA-B*39 allele increases type 1 diabetes risk conferred by HLA-DRB1*04:04-DQB1*03:02 and HLA-DRB1*08-DQB1*04 class II haplotypes. Hum Immunol 75, 65–70 (2014).

8. Study, T.I.H.C. The Major Genetic Determinants of HIV-1 Control Affect HLA Class I Peptide Presentation. Science 330, 1551–1557 (2010).

9. Lobos, C.A., Downing, J., D’Orsogna, L.J., Chatzileontiadou, D.S.M. & Gras, S. Protective HLA-B57: T cell and natural killer cell recognition in HIV infection. Biochemical Society Transactions 50, 1329–1339 (2022).

10. Ostrov, D.A. et al. Drug hypersensitivity caused by alteration of the MHC-presented self-peptide repertoire. Proc Natl Acad Sci U S A 109, 9959–64 (2012).

11. Kawase, T. et al. High-risk HLA allele mismatch combinations responsible for severe acute graft-versus-host disease and implication for its molecular mechanism. Blood 110, 2235–2241 (2007).

12. Marino, S.R. et al. Identification by random forest method of HLA class I amino acid substitutions associated with lower survival at day 100 in unrelated donor hematopoietic cell transplantation. Bone Marrow Transplantation 47, 217–226 (2012).

13. Illing, P.T. et al. HLA-B57 micropolymorphism defines the sequence and conformational breadth of the immunopeptidome. Nature Communications 9, 4693 (2018).

14. Fiorillo, M.T. et al. Allele-dependent Similarity between Viral and Self-peptide Presentation by HLA-B27 Subtypes*. Journal of Biological Chemistry 280, 2962–2971 (2005).

15. Macdonald, W.A. et al. A Naturally Selected Dimorphism within the HLA-B44 Supertype Alters Class I Structure, Peptide Repertoire, and T Cell Recognition. J. Exp. Med. 198, 679–691 (2003).

16. Saunders, P.M. et al. The molecular basis of how buried human leukocyte antigen polymorphism modulates natural killer cell function. Proc Natl Acad Sci U S A 117, 11636–11647 (2020).

17. Li, X. et al. Molecular basis of differential HLA class I-restricted T cell recognition of a highly networked HIV peptide. Nat Commun 14, 2929 (2023).

18. Kløverpris, H.N. et al. A molecular switch in immunodominant HIV-1-specific CD8 T-cell epitopes shapes differential HLA-restricted escape. Retrovirology 12, 20 (2015).

19. Luz, J.G. et al. Structural Comparison of Allogeneic and Syngeneic T Cell Receptor-Peptide-Major Histocompatibility Complex Complexes: A Buried Alloreactive Mutation Subtly Alters Peptide Presentation Substantially Increasing V{beta} Interactions. J. Exp. Med. 195, 1175–1186 (2002).

20. Zhang, S. et al. Structural basis of cross-allele presentation by HLA-A*0301 and HLA-A*1101 revealed by two HIV-derived peptide complexes. Molecular Immunology 49, 395–401 (2011).

21. Zhu, S. et al. Divergent Peptide Presentations of HLA-A*30 Alleles Revealed by Structures With Pathogen Peptides. Frontiers in Immunology **Volume** 10 - 2019(2019).

22. Archbold, J.K. et al. Natural micropolymorphism in human leukocyte antigens provides a basis for genetic control of antigen recognition. J. Exp. Med. 206, 209–219 (2009).

23. Sanderson, J.P. et al. Preclinical evaluation of an affinity-enhanced MAGE-A4-specific T-cell receptor for adoptive T-cell therapy. Oncoimmunology 9, 1682381 (2020).

24. Fabian, H. et al. HLA-B27 heavy chains distinguished by a micropolymorphism exhibit differential flexibility. Arthritis Rheum 62, 978–87 (2010).

25. Fabian, H. et al. HLA-B27 subtypes differentially associated with disease exhibit conformational differences in solution. Journal of Molecular Biology 376, 798–810 (2008).

26. Loll, B., Rückert, C., Uchanska-Ziegler, B. & Ziegler, A. Conformational Plasticity of HLA-B27 Molecules Correlates Inversely With Efficiency of Negative T Cell Selection. Frontiers in Immunology **Volume** 11 - 2020(2020).

27. Miles, J.J. et al. TCR alpha genes direct MHC restriction in the potent human T cell response to a class I-bound viral epitope. J Immunol 177, 6804–14 (2006).

28. Abualrous, E.T. et al. F pocket flexibility influences the tapasin dependence of two differentially disease-associated MHC Class I proteins. European Journal of Immunology 45, 1248–1257 (2015).

29. Bailey, A. et al. Selector function of MHC I molecules is determined by protein plasticity. Scientific Reports 5, 14928 (2015).

30. Amarajeewa, A.W.P. et al. Polymorphism in F pocket affects peptide selection and stability of type 1 diabetes-associated HLA-B39 allotypes. European Journal of Immunology 54, 2350683 (2024).

31. Serçinoğlu, O. & Ozbek, P. Computational characterization of residue couplings and micropolymorphism-induced changes in the dynamics of two differentially disease-associated human MHC class-I alleles. Journal of Biomolecular Structure and Dynamics 36, 724–740 (2018).

32. McShan, A.C. et al. Molecular determinants of chaperone interactions on MHC-I for folding and antigen repertoire selection. Proc Natl Acad Sci U S A 116, 25602–25613 (2019).

33. Chandran, S.S. et al. Immunogenicity and therapeutic targeting of a public neoantigen derived from mutated PIK3CA. Nat Med 28, 946–957 (2022).

34. Ma, J. et al. Dynamic allostery in the peptide/MHC complex enables TCR neoantigen selectivity. Nature Communications 16, 849 (2025).

35. Smith, A.R. et al. Structurally silent peptide anchor modifications allosterically modulate T cell recognition in a receptor-dependent manner. Proceedings of the National Academy of Sciences 118, e2018125118 (2021).

36. Miley, M.J. et al. Structural Basis for the Restoration of TCR Recognition of an MHC Allelic Variant by Peptide Secondary Anchor Substitution. J. Exp. Med. 200, 1445–1454 (2004).

37. Ding, Y.H., Baker, B.M., Garboczi, D.N., Biddison, W.E. & Wiley, D.C. Four A6-TCR/peptide/HLA-A2 structures that generate very different T cell signals are nearly identical. Immunity 11, 45–56 (1999).

38. Hawse, W.F. et al. Peptide Modulation of Class I Major Histocompatibility Complex Protein Molecular Flexibility and the Implications for Immune Recognition. Journal of Biological Chemistry 288, 24372–24381 (2013).

39. Hopkins, J.R. et al. Peptide cargo tunes a network of correlated motions in human leucocyte antigens. FEBS J 287, 3777–3793 (2020).

40. Mikhaylov, V. et al. Accurate modeling of peptide-MHC structures with AlphaFold. Structure 32, 228–241.e4 (2024).

41. Gupta, S., Nerli, S., Kutti Kandy, S., Mersky, G.L. & Sgourakis, N.G. HLA3DB: comprehensive annotation of peptide/HLA complexes enables blind structure prediction of T cell epitopes. Nature Communications 14, 6349 (2023).

42. Riley, T.P. et al. Structure Based Prediction of Neoantigen Immunogenicity. Frontiers in Immunology 10(2019).

43. Antunes, D.A. et al. Interpreting T-Cell Cross-reactivity through Structure: Implications for TCR-Based Cancer Immunotherapy. Frontiers in Immunology 8(2017).

44. Gonzalez-Galarza, Faviel F. et al. Allele frequency net database (AFND) 2020 update: gold-standard data classification, open access genotype data and new query tools. Nucleic Acids Research 48, D783–D788 (2019).

45. Singh, N.K. et al. An Engineered T Cell Receptor Variant Realizes the Limits of Functional Binding Modes. Biochemistry 59, 4163–4175 (2020).

46. Morgan, C.S., Holton, J.M., Olafson, B.D., Bjorkman, P.J. & Mayo, S.L. Circular dichroism determination of class I MHC-peptide equilibrium dissociation constants. Protein Sci 6, 1771–3 (1997).

47. Hellman, L.M. et al. Differential scanning fluorimetry based assessments of the thermal and kinetic stability of peptide-MHC complexes. J Immunol Methods 432, 95–101 (2016).

48. Ayres, C.M. & Baker, B.M. Peptide-dependent tuning of major histocompatibility complex motional properties and the consequences for cellular immunity. Current Opinion in Immunology 76, 102184 (2022).

49. Kutzner, C., Czub, J. & Grubmüller, H. Keep It Flexible: Driving Macromolecular Rotary Motions in Atomistic Simulations with GROMACS. J Chem Theory Comput 7, 1381–1393 (2011).

50. Nguyen, A.T. et al. Homologous peptides derived from influenza A, B and C viruses induce variable CD8+ T cell responses with cross-reactive potential. Clinical & Translational Immunology 11, e1422 (2022).

51. Sim, M.J.W. et al. Identification and structural characterization of a mutant KRAS-G12V specific TCR restricted by HLA-A3. European Journal of Immunology 54, 2451079 (2024).

52. Li, L. & Bouvier, M. Structures of HLA-A*1101 Complexed with Immunodominant Nonamer and Decamer HIV-1 Epitopes Clearly Reveal the Presence of a Middle, Secondary Anchor Residue1. The Journal of Immunology 172, 6175–6184 (2004).

53. Liu, W.J. et al. Cross-immunity Against Avian Influenza A(H7N9) Virus in the Healthy Population Is Affected by Antigenicity-Dependent Substitutions. The Journal of Infectious Diseases 214, 1937–1946 (2016).

54. Nguyen, A.T. et al. SARS-CoV-2 Spike-Derived Peptides Presented by HLA Molecules. Biophysica 1, 194–203 (2021).

55. Poole, A. et al. Therapeutic high affinity T cell receptor targeting a KRASG12D cancer neoantigen. Nature Communications 13, 5333 (2022).

56. Habel, J.R. et al. HLA-A*11:01-restricted CD8+ T cell immunity against influenza A and influenza B viruses in Indigenous and non-Indigenous people. PLOS Pathogens 18, e1010337 (2022).

57. Ahn, Y.M. et al. The impact of SARS-CoV-2 spike mutation on peptide presentation is HLA allomorph-specific. Current Research in Structural Biology 7, 100148 (2024).

58. Kostyu, D.D. et al. HLA class I polymorphism: structure and function and still questions. Hum Immunol 57, 1–18 (1997).

59. Serçinoğlu, O. & Ozbek, P. Sequence-structure-function relationships in class I MHC: A local frustration perspective. PLoS One 15, e0232849 (2020).

60. Tussey, L.G. et al. Analysis of mutant HLA-A2 molecules. Differential effects on peptide binding and CTL recognition. J Immunol 152, 1213–21 (1994).

61. Mattson, D.H. et al. Differential effects of amino acid substitutions in the beta-sheet floor and alpha-2 helix of HLA-A2 on recognition by alloreactive viral peptide-specific cytotoxic T lymphocytes. J Immunol 143, 1101–7 (1989).

62. Matsui, M., Hioe, C. & Frelinger, J. Roles of the Six Peptide-Binding Pockets of the HLA-A2 Molecule in Allorecognition by Human Cytotoxic T-Cell Clones. PNAS 90, 674–678 (1993).

63. Jiang, X. & Fares, M.A. IDENTIFYING COEVOLUTIONARY PATTERNS IN HUMAN LEUKOCYTE ANTIGEN (HLA) MOLECULES. Evolution 64, 1429–1445 (2010).

64. Truong, H.V. & Sgourakis, N.G. Dynamics of MHC-I molecules in the antigen processing and presentation pathway. Curr Opin Immunol 70, 122–128 (2021).

65. Ilca, F.T., Drexhage, L.Z., Brewin, G., Peacock, S. & Boyle, L.H. Distinct Polymorphisms in HLA Class I Molecules Govern Their Susceptibility to Peptide Editing by TAPBPR. Cell Reports 29, 1621–1632.e3 (2019).

66. Badrinath, S., Kunze-Schumacher, H., Blasczyk, R., Huyton, T. & Bade-Doeding, C. A Micropolymorphism Altering the Residue Triad 97/114/156 Determines the Relative Levels of Tapasin Independence and Distinct Peptide Profiles for HLA-A(*)24 Allotypes. J Immunol Res 2014, 298145 (2014).

67. Thomas, C. & Tampé, R. MHC I assembly and peptide editing — chaperones, clients, and molecular plasticity in immunity. Current Opinion in Immunology 70, 48–56 (2021).

68. Thomas, C. & Tampé, R. Proofreading of Peptide—MHC Complexes through Dynamic Multivalent Interactions. Frontiers in Immunology 8, 65 (2017).

69. McMaster, B., Thorpe, C.J., Rossjohn, J., Deane, C.M. & Koohy, H. Quantifying conformational changes in the TCR:pMHC-I binding interface. Frontiers in Immunology **Volume** 15 - 2024(2024).

70. Ayres, C.M. et al. Dynamically Driven Allostery in MHC Proteins: Peptide-Dependent Tuning of Class I MHC Global Flexibility. Frontiers in Immunology 10, 1–13 (2019).

71. Ayres, C.M., Corcelli, S.A. & Baker, B.M. Peptide and Peptide-Dependent Motions in MHC Proteins: Immunological Implications and Biophysical Underpinnings. Frontiers in Immunology 8, 1–9 (2017).

72. Smock, R.G. & Gierasch, L.M. Sending Signals Dynamically. Science 324, 198–203 (2009).

73. Cooper, A. & Dryden, D.T.F. Allostery without conformational change. European Biophysics Journal 11, 103–109 (1984).

74. Nussinov, R. Introduction to Protein Ensembles and Allostery. Chemical Reviews 116, 6263–6266 (2016).

75. Ma, J., Ayres, C.M., Hellman, L.M., Devlin, J.R. & Baker, B.M. Dynamic allostery controls the peptide sensitivity of the Ly49C natural killer receptor. Journal of Biological Chemistry 296(2021).

76. Feng, Y., Reinherz, E.L. & Lang, M.J. αβ T Cell Receptor Mechanosensing Forces out Serial Engagement. Trends Immunol 39, 596–609 (2018).

77. Zhao, X. et al. Tuning T cell receptor sensitivity through catch bond engineering. Science 376, eabl5282 (2022).

78. Liu, B., Chen, W., Evavold, Brian D. & Zhu, C. Accumulation of Dynamic Catch Bonds between TCR and Agonist Peptide-MHC Triggers T Cell Signaling. Cell 157, 357–368 (2014).

79. Sibener, L.V. et al. Isolation of a Structural Mechanism for Uncoupling T Cell Receptor Signaling from Peptide-MHC Binding. Cell 174, 672–687 e27 (2018).

80. Brazin, K.N. et al. Structural Features of the αβTCR Mechanotransduction Apparatus That Promote pMHC Discrimination. Front Immunol 6, 441 (2015).

81. Ayres, C.M., Corcelli, S.A. & Baker, B.M. The Energetic Landscape of Catch Bonds in TCR Interfaces. J Immunol 211, 325–332 (2023).

82. Otwinowski, Z. & Minor, W. Processing of X-ray Diffraction Data Collected in Oscillation Mode. Methods in Enzymology 276, 307–326 (1997).

83. Afonine, P.V. et al. Towards automated crystallographic structure refinement with phenix.refine. Acta Crystallographica Section D 68, 352–367 (2012).

84. Agirre, J. et al. The CCP4 suite: integrative software for macromolecular crystallography. Acta Crystallographica Section D 79, 449–461 (2023).

85. Emsley, P., Lohkamp, B., Scott, W.G. & Cowtan, K. Features and development of Coot. Acta Crystallographica Section D 66, 486–501 (2010).

86. Hsin, J., Arkhipov, A., Yin, Y., Stone, J.E. & Schulten, K. Using VMD: an introductory tutorial. Curr Protoc Bioinformatics **Chapter** 5, Unit 5.7 (2008).

87. Pettersen, E.F. et al. UCSF Chimera—A visualization system for exploratory research and analysis. Journal of Computational Chemistry 25, 1605–1612 (2004).

88. Jurcik, A. et al. CAVER Analyst 2.0: analysis and visualization of channels and tunnels in protein structures and molecular dynamics trajectories. Bioinformatics 34, 3586–3588 (2018).

89. Blevins, S.J. & Baker, B.M. Using Global Analysis to Extend the Accuracy and Precision of Binding Measurements with T cell Receptors and Their Peptide/MHC Ligands. Frontiers in Molecular Biosciences 4, 1–9 (2017).

90. Yi, F. et al. CAR-engineered lymphocyte persistence is governed by a FAS ligand–FAS autoregulatory circuit. Nature Cancer 6, 1638–1655 (2025).

91. Roe, D.R. & Cheatham, T.E. PTRAJ and CPPTRAJ: Software for Processing and Analysis of Molecular Dynamics Trajectory Data. Journal of Chemical Theory and Computation 9, 3084–3095 (2013).

