## Supplemental figures and table for "HLA micropolymorphisms confine neoantigen conformational adaptability and guide T cell receptor selectivity"

### FIGURE S1

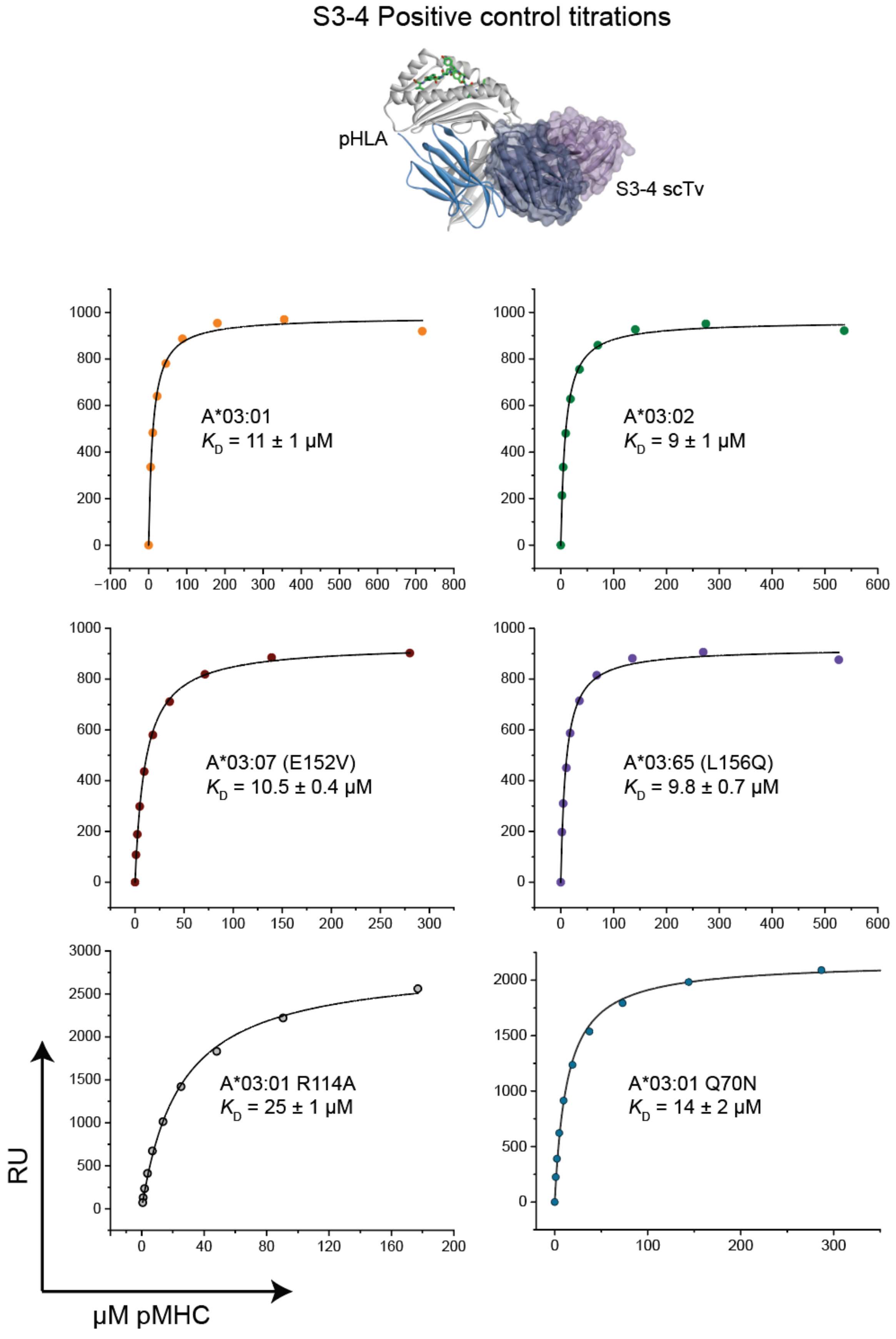

S3-4 Positive control titrations
pHLA
S3-4 scTv
A*03:01
$K_D$ = 11 ± 1 μM
A*03:02
$K_D$ = 9 ± 1 μM
A*03:07 (E152V)
$K_D$ = 10.5 ± 0.4 μM
A*03:65 (L156Q)
$K_D$ = 9.8 ± 0.7 μM
A*03:01 R114A
$K_D$ = 25 ± 1 μM
A*03:01 Q70N
$K_D$ = 14 ± 2 μM
RU
μM pMHC

### FIGURE S2

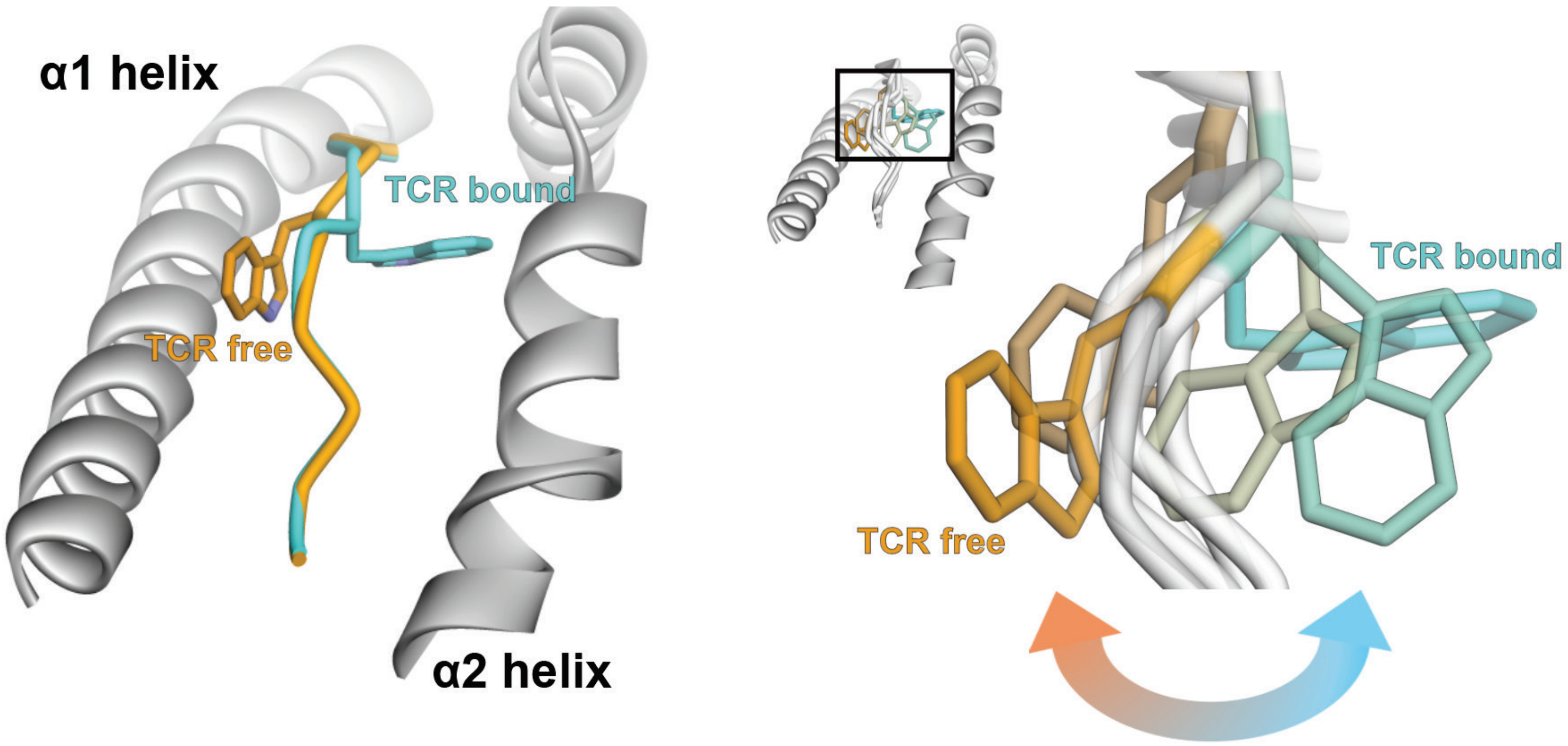

**FIGURE S3**

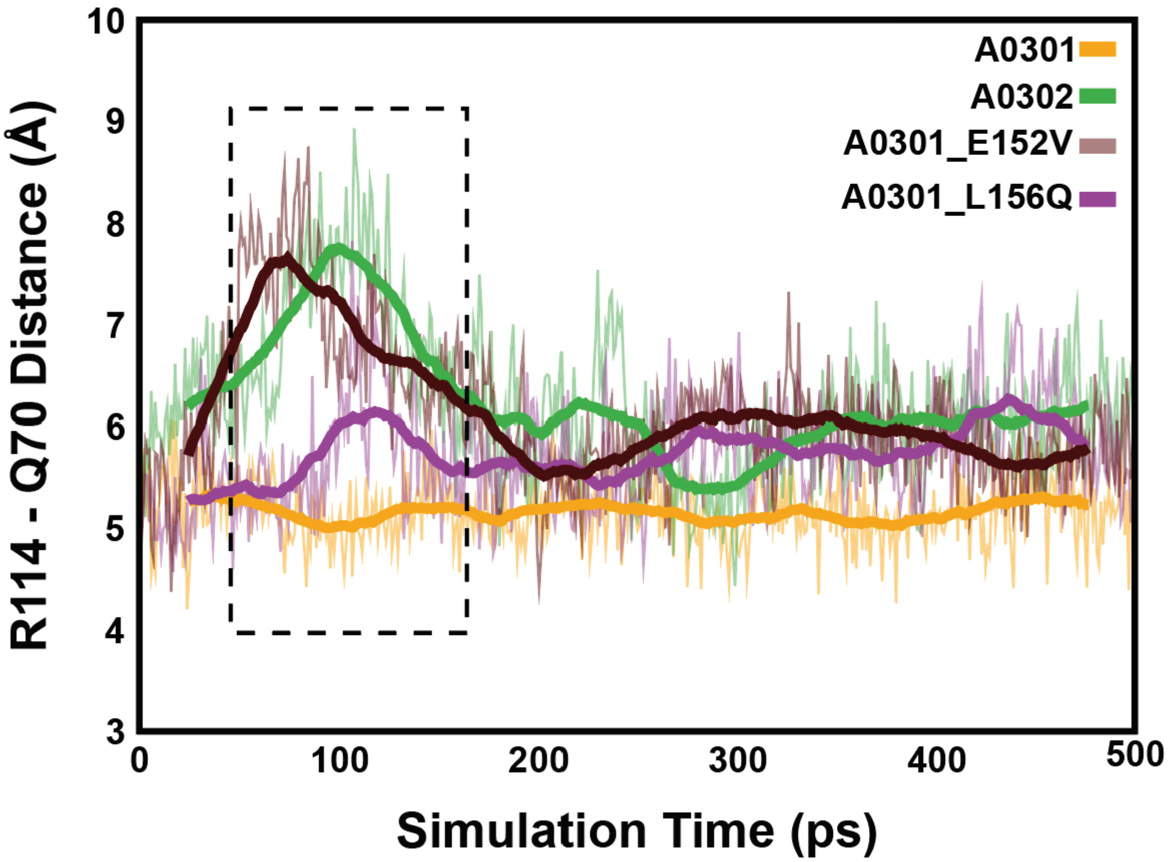

A0301
A0302
A0301_E152V
A0301_L156Q
R114 - Q70 Distance (Å)
Simulation Time (ps)
10
9
8
7
6
5
4
3
0
100
200
300
400
500

### TABLE S1

| | **neoAg/HLA-A*03:02** | **neoAg/HLA-A*03:07 (E152V)** | **neoAg/HLA-A*03:65 (L156Q)** | **neoAg/HLA-A*03:65 (L156Q; alternate)** |
|---|---|---|---|---|
| **Data Collection** | | | | |
| Space group | P 6 2 2 | P 6 2 2 | P 6 2 2 | P 6 2 2 |
| Unit cell dimensions | | | | |
| *a, b, c* (Å) | 156.03, 156.03, 85.58 | 156.06, 156.06, 85.51 | 156.09, 156.09, 85.78 | 156.42, 156.42, 85.63 |
| *α, β, γ* (°) | 90, 90, 120 | 90, 90, 120 | 90, 90, 120 | 90, 90, 120 |
| Resolution (Å) | 50.00-2.05 (2.09-2.05) | 50.00-1.92 (1.95-1.92) | 50.0-2.00 (2.03-2.00) | 50.00-2.30 (2.30-2.34) |
| $R_{merge}$ | 0.15 (1.77) | 0.15 (1.11) | 0.20 (0.75) | 0.10 (0.76) |
| $I/\sigma I$ | 45.94 (4.43) | 42.90 (4.9) | 23.14 (5) | 50.78 (9.2) |
| Completeness (%) | 96.9 (79.0) | 97.4 (85.8) | 97.9 (82.5) | 98.94 (96.19) |
| Total reflections | 6182506 | 9675781 | 9126650 | 1103862 |
| Unique Reflections | 37920 (1110) | 45911 (1333) | 41337 (1162) | 27935 (2628) |
| Redundancy | 37.4 (34.3) | 30.4 (20.3) | 36.5 (25.6) | 39.5 (36.3) |
| **Refinement** | | | | |
| Resolution (Å) | 45.04-2.05 | 45.05-1.92 | 43.9-2.00 | 39.94-2.30 |
| $R_{work}/R_{free}$ | 0.21/0.24 | 0.21/0.24 | 0.20/0.22 | 0.19/0.22 |
| Number of atoms | | | | |
| Total | 3517 | 3586 | 3520 | 3435 |
| Protein | 3126 | 3125 | 3135 | 3139 |
| Average B-factors (Å) | 38.39 | 30.92 | 34.15 | 45.02 |
| R.M.S deviations | | | | |
| Bond length (Å) | 0.004 | 0.005 | 0.003 | 0.002 |
| Bond angles (°) | 0.61 | 0.7 | 0.62 | 0.44 |
| Ramachandran favored (%) | 98.4 | 97.87 | 98.14 | 97.08 |
| Ramachandran outliers (%) | 0 | 0 | 0 | 0 |
| PDB accession code | 9CWZ | 9CX0 | 9CX1 | 9CX2 |
